# Multiomic Single Cell Sequencing Identifies Stemlike Nature of Mixed Phenotype Acute Leukemia and Provides Novel Risk Stratification

**DOI:** 10.1101/2023.05.15.540305

**Authors:** Cheryl A. C. Peretz, Vanessa E. Kennedy, Anushka Walia, Cyrille L. Delley, Andrew Koh, Elaine Tran, Iain C. Clark, Corey E. Hayford, Chris D’Amato, Yi Xue, Kristina M. Fontanez, Ritu Roy, Aaron C. Logan, Alexander E. Perl, Adam Abate, Adam Olshen, Catherine C. Smith

**Affiliations:** Divison of Hematology and Oncology, Department of Pediatrics, USA; Helen Diller Family Comprehensive Cancer Center, University of California San Francisco, San Francisco, CA; Division of Hematology and Oncology, Department of Medicine, University of California San Francisco, San Francisco, CA, USA; Bioengineering and Therapeutic Sciences, University of California San Francisco, San Francisco, CA, USA; Bioengineering, University of California Berkeley, Berkeley, CA, USA; Fluent Biosciences Inc., Watertown, MAw; Department of Epidemiology and Biostatistics, Universiwty of California San Francisco, San Francisco, CA, USA; Department of Medicine, Division of Hematology-Oncology, Perelman School of Medicine at the University of Pennsylvania, Philadelphia, PA, USA

## Abstract

Mixed phenotype acute leukemia (MPAL) is a leukemia whose biologic drivers are poorly understood, therapeutic strategy remains unclear, and prognosis is poor. We performed multiomic single cell (SC) profiling of 14 newly diagnosed adult MPAL patients to characterize the immunophenotypic, genetic, and transcriptional landscapes of MPAL. We show that neither genetic profile nor transcriptome reliably correlate with specific MPAL immunophenotypes. However, progressive acquisition of mutations is associated with increased expression of immunophenotypic markers of immaturity. Using SC transcriptional profiling, we find that MPAL blasts express a stem cell-like transcriptional profile distinct from other acute leukemias and indicative of high differentiation potential. Further, patients with the highest differentiation potential demonstrated inferior survival in our dataset. A gene set score, MPAL95, derived from genes highly enriched in this cohort, is applicable to bulk RNA sequencing data and was predictive of survival in an independent patient cohort, suggesting utility for clinical risk stratification.

## Introduction

Mixed phenotype acute leukemia (MPAL) is characterized by leukemic blasts with both lymphoid and myeloid cell surface markers. Survival of patients with MPAL is poor and inferior to that of the more common acute lymphoid and myeloid leukemias (ALL and AML)^1^. The diagnostic definition of MPAL remains unrefined. While both ALL and AML are defined by genetic drivers, the 2022 WHO guidelines define MPAL largely by immunophenotype and include of only a handful of defining genetic abnormalities (*BCR*::*ABL1* fusion, *KMT2A, ZNF384, and BCL11B* rearrangements)^2^. Genomic alterations in MPAL are not unique and include mutations recurrently mutated in ALL or AML^3^. The biologic connection between immunophenotype and genotype in MPAL is unknown. Importantly, neither the immunophenotype nor the genotype of MPAL correlates with overall survival, suggesting that a more complete biologic understanding of MPAL, and subsequent disease definition and risk stratification, remains to be determined^2, 4^.

Due to the relative rarity and heterogeneous nature of MPAL, optimal therapeutic strategies remain uncertain. Emerging data suggests that sub-classification of MPAL may be needed to facilitate therapeutic decision making^5^. However, the full immunophenotypic, genetic, and transcriptomic changes that determine risk stratification of this complex disease have not been elucidated. Until recently, the technology to simultaneously determine immunophenotypic, genetic, and transcriptomic heterogeneity in MPAL has not existed. MPAL, with its definitionally ‘mixed’ immunophenotype, is uniquely poised to benefit from multiomic single cell (SC) sequencing analysis, which can, for the first time, quantify the relationship between these cellular factors to better understand the biologic origin of MPAL and the mechanism of its poor prognosis.

Here, we use multiomic SC profiling on newly diagnosed MPAL samples to characterize the immunophenotypic, genetic, and transcriptional landscapes of MPAL. This profiling allows us to contradict the paradigm of MPAL as an entity on a continuum with ALL and AML, and instead identify MPAL as a distinct, stem-like disease that, in contrast to other leukemias, cannot be defined by genetics alone. We further describe a stem cell-derived gene expression signature for MPAL that can predict patient survival. These results broaden our understanding of MPAL biology and suggest a path toward novel risk stratification. Importantly, these findings have potential to direct treatment and, hopefully, ultimately improve the prognosis of this disease.

## Results

### The Heterogeneous Genetic Landscape of MPAL

To characterize the heterogeneous genetic, transcriptional, and immunophenotypic landscape of MPAL, we analyzed samples from 14 patients with newly diagnosed MPAL using two SC technologies in parallel: DAb-Seq (SC DNA plus protein sequencing)^6–8^ and CITE-seq (SC RNA plus protein sequencing)^9^ (Figure 1A). Patient characteristics are in Supplementary Table 1. By clinical immunophenotyping via flow cytometry, our cohort included 10 patients with B/myeloid, 3 patients with T/myeloid, and 1 patient with B and T/myeloid MPAL.

**Figure 1:**
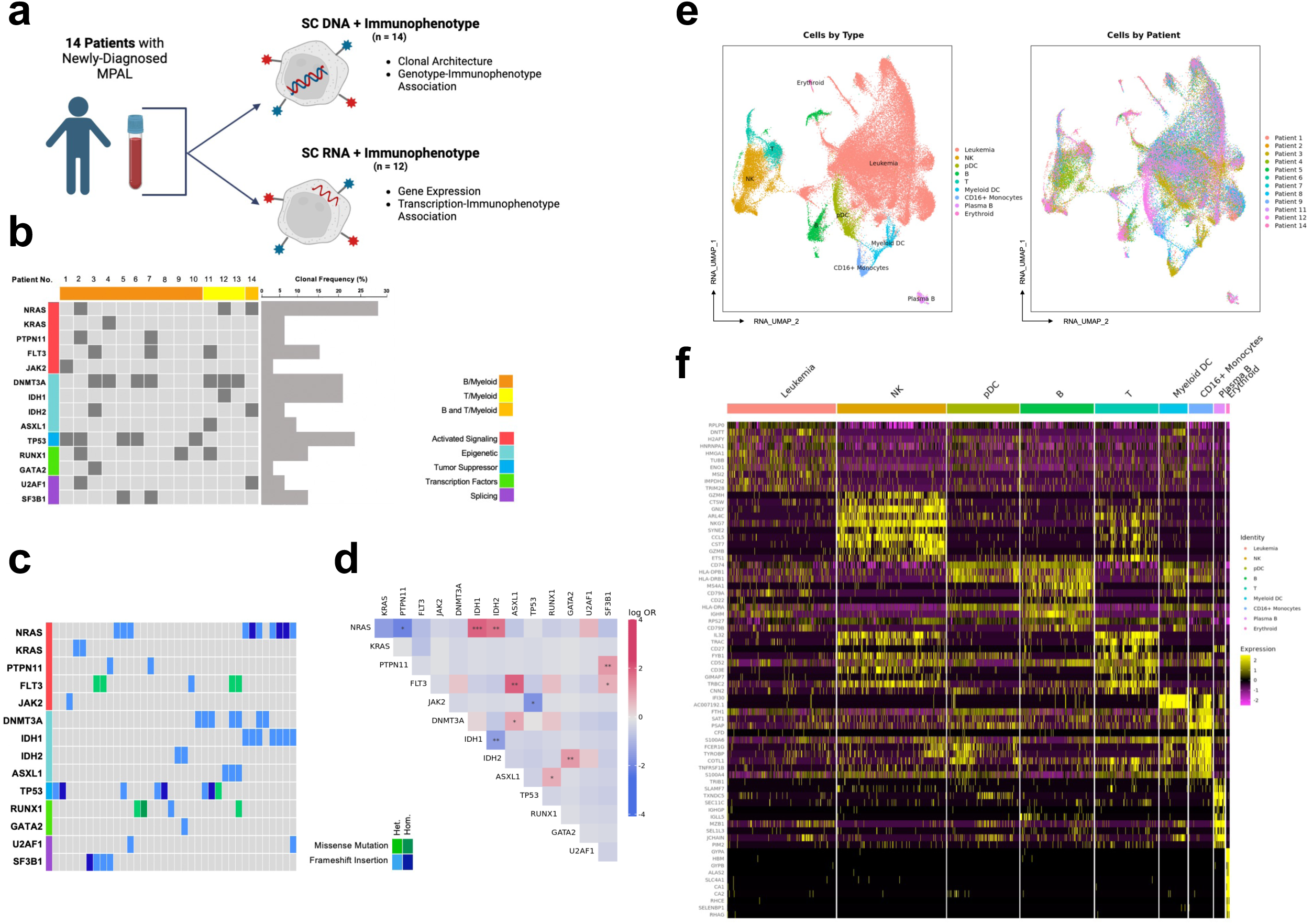
Single-cell (SC) genetic, immunophenotypic, and transcriptomic landscape of mixed phenotypic acute leukemia (MPAL). **A.** Schematic depicting sample workflow. **B.** Oncoprint of 14 patients with newly diagnosed MPAL. Each column is a unique patient. Patients are coded on the top row based on immunophenotypic subtype and mutations are ordered based on biologic function. Clonal frequency is based on the total number of clones the mutation was present in, not accounting for zygosity. **C.** Oncoprint of 36 genetically defined clones across 14 patients with MPAL. Each column is a unique clone, and mutations are color-coded based on type of mutation and zygosity. **D.** Pairwise association of driver mutations identified via SC DNA sequencing across 36 clones in 14 patients with MPAL. For each mutation pair, cooccurrence is summarized as log odds ratio (OR), with positive values indicating cooccurrence and negative values mutual exclusivity. Statistical significance is indicated as *p, .05; **p, .01; ***p, .001. **E.** RNA-derived UMAP from SC CITE-seq analysis of 71,579 cells from 12 patients. Cells are color-coded by cell lineage/type as determined by gene expression data (*left*) and by individual patient (*right*). **F.** Heatmap of scaled expression values for top ten most upregulated conserved genes for each transcriptionally defined cell type as identified in **1E**.

For DAb-seq, we used a panel covering hotspots in 20 genes frequently mutated in leukemia combined with 22 antibody-oligonucleotide conjugates for cell-surface immunophenotypic proteins on hematopoietic cells (Supplementary Table 2, Supplementary Table 3)^6–8^. A total of 58,807 individual cells were genotyped, with a median of 4,221 cells per sample (range 1,093 - 7,245 cells/sample) (Supplementary Table 4).

The mutational landscape for all patients and clones is depicted in Figures 1B and 1C. Across the cohort, we identified 27 pathogenic or likely pathogenic mutations within 36 genetically-distinct clones (median 2.6 clones/patient, range 0 - 6); there was no difference in the number of clones between B/myeloid and T/myeloid MPAL (2.8 vs 2.3, p = 0.66) (Supplementary Table 5). At the clone level, the most commonly mutated genes were *NRAS*, present in 10 clones (28%), *TP53*, present in 8 clones (22%), and *DNMT3A* and *IDH1*, each present in 7 clones (19%).

Clone-level mutational co-occurrence analysis demonstrated the strongest positive association between *NRAS*/*IDH1* (Odds Ratio [OR] 8.91, p <0.0001), *FLT3/ASXL1* (OR 8.58, p = 0.008) and *PTPN11/SF3B1* (OR 4.13, p = 0.002); *IDH1/IDH2* were negatively associated (OR -0.58, p = 0.003) (Figure 1D). Except for *DNMT3A/ASXL1*, mutations from the same functional class were infrequently co-mutated in the same single cell and clone; notably, no clones demonstrated more than one distinct signaling mutation.

Using SC DNA sequencing, we reconstructed the evolutionary history of each patient using single cell inference of tumor evolution (SCITE), a probabilistic model to infer genetic phylogeny (Supplementary Figure 1)^10^. Patients demonstrated diverse phylogenetic trees with both linear and branched architectures. Across the cohort, the most common functional class of founding mutations was epigenetic regulators, at 7/18 (38.8%). The most common functional class of branch mutations was activated signaling mutations, at 10/25 (40%).

### The Heterogeneous Transcriptional Landscape of MPAL

For CITE-seq analysis, we used a particle-templated instant partitions sequencing (PIPseq) approach to perform SC indexing of transcriptomes and epitomes sequencing (CITE-seq) analysis with a panel of 19 barcoded antibodies (Supplementary Table 6)^9^. A total of 72,131 individual cells from 12 patients were genotyped, with a median of 6,010 cells per sample (range 1,173 – 10,275) (Supplementary Table 4). Across all patients, SC transcriptional data was clustered by transcription and annotated (Figure 1E). Notably, all 12 patients, regardless of MPAL immunophenotypic subtype, contributed to the cluster annotated as leukemia (Supplementary Figure 2). After normalization for the number of cells isolated per patient, the proportion of the leukemia cluster attributed to an individual patient ranged from 4.5% - 10.4% (median 8.8%). Furthermore, immunophenotypic subtype was not the primary predictor of transcriptional variation in correspondence analysis (Supplementary Figure 3). Relative to non-leukemic cells and clusters, the conserved leukemia cluster demonstrated a unique transcriptional signature (Figure 1F, Supplementary Table 7).

### Genotype and Immunophenotype Incompletely Associate Across Patients

Using DAb-seq, we examined the association between immunophenotype and genetic clonal architecture. Patients with MPAL demonstrated heterogeneous immunophenotypes among both individual patients and MPAL subtypes (Supplementary Figure 4A, B). Across all patients, we observed some broad genotype-immunophenotype associations, including associations between *JAK2* mutations and CD71 (Point-biserial correlation coefficient 0.98; p < 7.2e-8), *NRAS* and CD38 (Point-biserial correlation coefficient 0.89; p = 0.004), and *IDH2* and CD11b and CD64 (Point-biserial correlation coefficients 0.87 and 0.80; p=0.002 and p=0.008, respectively) (Figure 2A, Supplementary Figure 4C).

**Figure 2:**
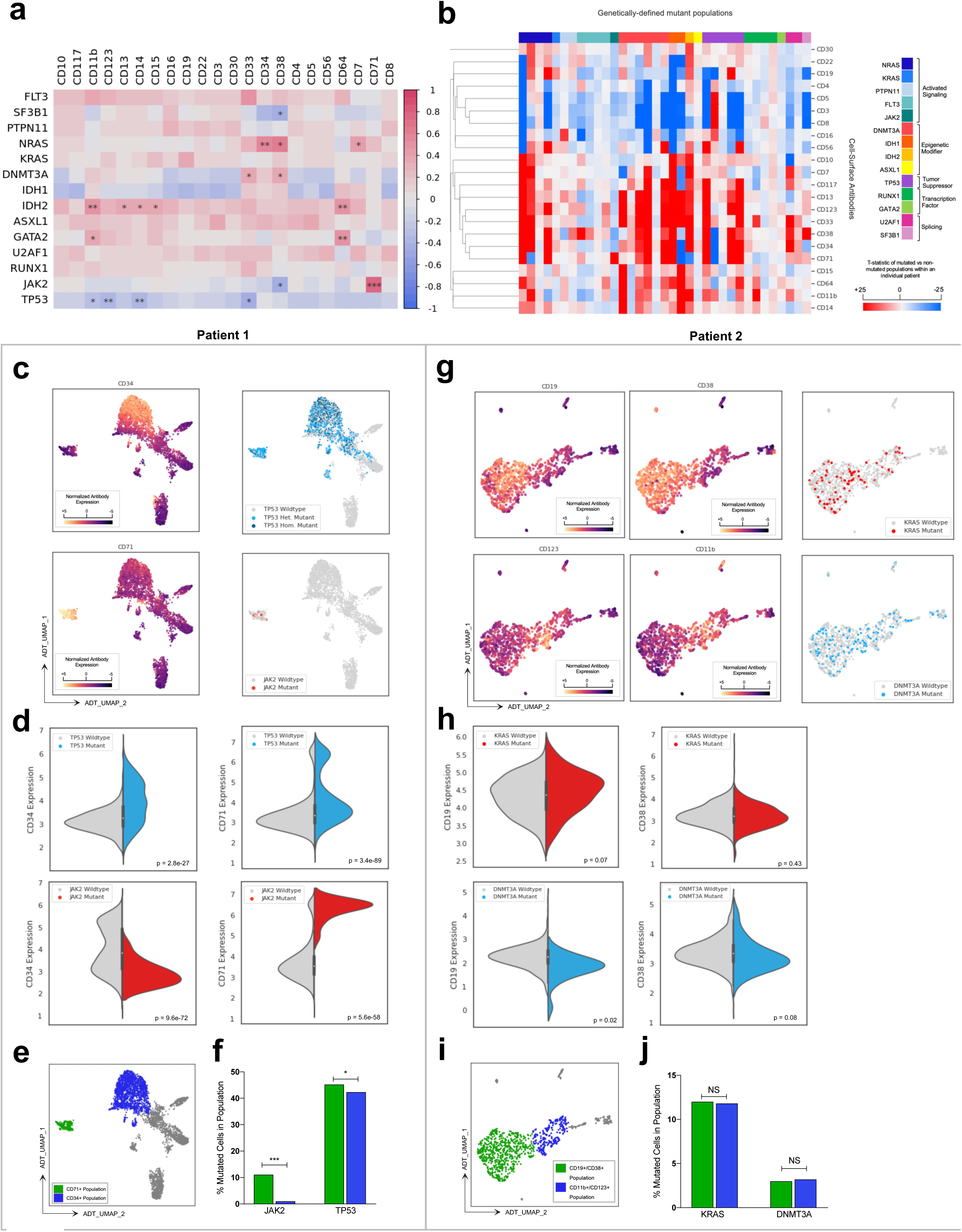
Heterogeneous genotype-immunophenotype association in MPAL. For panels C- J, each column represents a unique patient (*Left:* Patient 1; *Right:* Patient 2). **A.** Spearman correlation matrix across 36 unique genetically defied clones and 22 cell-surface antibodies. Correlation coefficient is denoted by color coding from highly correlated (red) to highly anti-correlated (blue), with significance denoted as *p, .05; **p, .01; ***p, .001. **B.** Heatmap of T-statistics generated by comparing cell-surface antibody expression of mutant vs non-mutant cell populations within an individual patient. Comparisons are only made within individual patients, not across patients. Rows are cell-surface antibodies with unsupervised hierarchical clustering applied. Columns are mutated populations color-coded by gene and ordered by biologic function. **C.** Immunophenotype-derived UMAP of 4,274 cells from **Patient 1**. Cells are color-coded based on CD34 expression (*top left*), CD71 expression (*bottom left)*, presence of TP53 mutations (*top right)* and presence of JAK2 mutations (*bottom right*). **D.** Violin plot comparing expression of CD34 and CD71 for TP53-mutated vs TP53-wildtype (WT) cells (*top)* and JAK2-mutated vs JAK2-WT cells (*bottom*) for **Patient 1**. The grey half of the split-violin splot represents non-mutated cells and the colorful half of the plot represents mutated cells within the same patient. **E.** UMAP from **2C** with unsupervised hierarchical clustering of cells into subpopulations manually annotated by immunophenotype. Clustering identified distinct CD34+ and CD71+ cell populations. **F.** Bar graph comparing the percentage of JAK2-mutated and TP53-mutated cells in the immunophenotypically-annotated populations identified in **2E**. **G.** Immunophenotype-derived UMAP of 2,848 cells from **Patient 2**. Cells are color-coded based on CD19 (*top left)*, CD38 (*top center*), CD11b (*bottom left*), and CD123 (*bottom center*) expression and presence of KRAS and DNMT3A mutations (*top right, bottom right)*. **H.** Violin plot comparing expression of CD19 and CD38 for KRAS-mutated vs KRAS-WT cells (*top*) and DNMT3A-mutated vs DNMT3A-WT cells *(bottom*) for **Patient 2**. **I.** UMAP from **2G** with unsupervised hierarchical clustering of cells into subpopulations manually annotated by immunophenotype. Clustering identified distinct CD19+/CD38+ and CD11b+/CD123+ cell populations. **J.** Bar graph comparing the percentage of KRAS-mutated and DNMT3A-mutated cells in the immunophenotypically-annotated populations identified in **2I.** Statistical significance is denoted as *p, .05; **p, .01; ***p, .001, with all p-values adjusted via the Bonferroni method.

We also observed considerable inter- and intra-patient heterogeneity at the clonal level; genotype-immunophenotype associations were present in some, but not all, mutated clonal populations (Figure 2B). For instance, our cohort included 4 *NRAS*-mutated clones. In 3/4, *NRAS*-mutated cells had significantly increased CD34 expression relative to *NRAS*-wildtype (WT) blasts within the same patient (t-statistics 52.3, 20.1, 22.3; p = 0.0, p = 1.7e-85, p = 3e- 99); however, in one clone there was no difference in CD34 expression between *NRAS*-mutated vs *NRAS-*WT cells (t-statistic 1.2; p = 0.25). Increased expression of other immunophenotypic proteins associated with an immature cell state, including CD38, CD33, CD123, and CD117, was also observed among select *NRAS*-mutated populations. These striking relationships were not observed, however, among other mutations in genes associated with cell signaling, including *KRAS*, *PTPN11*, or *FLT3* (Supplementary Figure 5A). Similarly, select *DNMT3A, IDH1* and *IDH2* mutated populations were associated with markedly increased expression of CD13 and CD11b, both associated with myeloid/monocytic differentiation, but this pattern was not consistent among all clones with these mutations (Supplementary Figure 5B).

### Genotype and Immunophenotype Incompletely Associate Within Individual Patients

The heterogeneous association between genotype and immunophenotype was further observed at the individual patient level. In some patients, genotype was strikingly associated with a distinct immunophenotype, while in other patients, genotype-immunophenotype association was not observed. For example, in Patient 1, we identified multiple immunophenotypically defined subpopulations, including a subset of cells with strong CD34 expression and a subset with strong CD71 expression (Figure 2C). In this patient, *JAK2-*mutated cells were associated with significantly higher CD71 expression relatively to *JAK2-*WT cells (median CD71 expression 6.40 vs 3.49, p =3.4e-89), and the proportion of *JAK2-*mutated cells in the CD71-bright population was higher than in the CD34-bright population (11.1% vs 0%, p < 1e-99) (Figure 2D-F). By contrast, in Patient 2, genotype and immunophenotype were not associated. Like Patient 1, Patient 2 also had multiple immunophenotypically-defined subpopulations, including a population with strong CD19 and CD38 expression and a separate population with strong CD11b and CD123 expression (Figure 2G). Unlike Patient 1, in Patient 2, identified driver mutations were not associated with a specific immunophenotype, with an equal proportion of *KRAS*-mutated and *DNMT3A-*mutated cells in the CD19+/CD38+ population vs the CD11b+/CD123+ population (*KRAS:* 12.01 vs. 11.84%, p = 0.06; *DNMT3A:* 3.08 vs 3.19%, p =0.22) (Figure 2H-J).

Further highlighting the heterogeneity among genotype-immunophenotype associations, we observed that the same mutation does not consistently associate with the same immunophenotype across patients. For example, both Patient 7 and Patient 14 harbor an *IDH2* R140Q mutation. In Patient 7, *IDH2-*mutated cells were significantly associated with increased expression of monocytic markers CD11b, CD64, CD13, and CD14 relative to *IDH2-*WT cells (median CD11b expression 4.12 vs 5.54, p = 9e-88; CD64 2.01 vs 2.89, p = 1.3e-34; CD13 3.34 vs 4.75, p = 2.3e-58; CD14 3.38 vs 3.90, p = 8.8e-40) (Supplementary Figure 6A-B). Although Patient 14 had the same *IDH2* R140Q mutation, *IDH2-*mutated cells in this patient only demonstrated slightly higher expression of CD11b and did not have higher expression of other monocytic markers (median CD11b expression 3.29 vs 3.67, p = 0.012; CD64 1.04 vs 1.11, p = 0.12; CD13 3.06 vs 3.20, p = 0.09; CD14 2.76 vs 2.88, p = 0.21) (Supplementary Figure 6).

### Progressive Mutational Acquisition is Associated with Increase in Expression of Immunophenotypic Markers of Immaturity

In addition to the association between genotype and immunophenotype, we also assessed the association between mutational phylogenetic progression and immunophenotypic evolution. Of the 14 patients in our cohort, 9 had at least 2 stepwise mutational acquisitions identified on SC phylogenetic analysis (Supplementary Figure 1). For these 9 patients, we measured how cell-surface immunophenotypic protein expression changed with progressive acquisition of mutations (Figure 3A).

**Figure 3.**
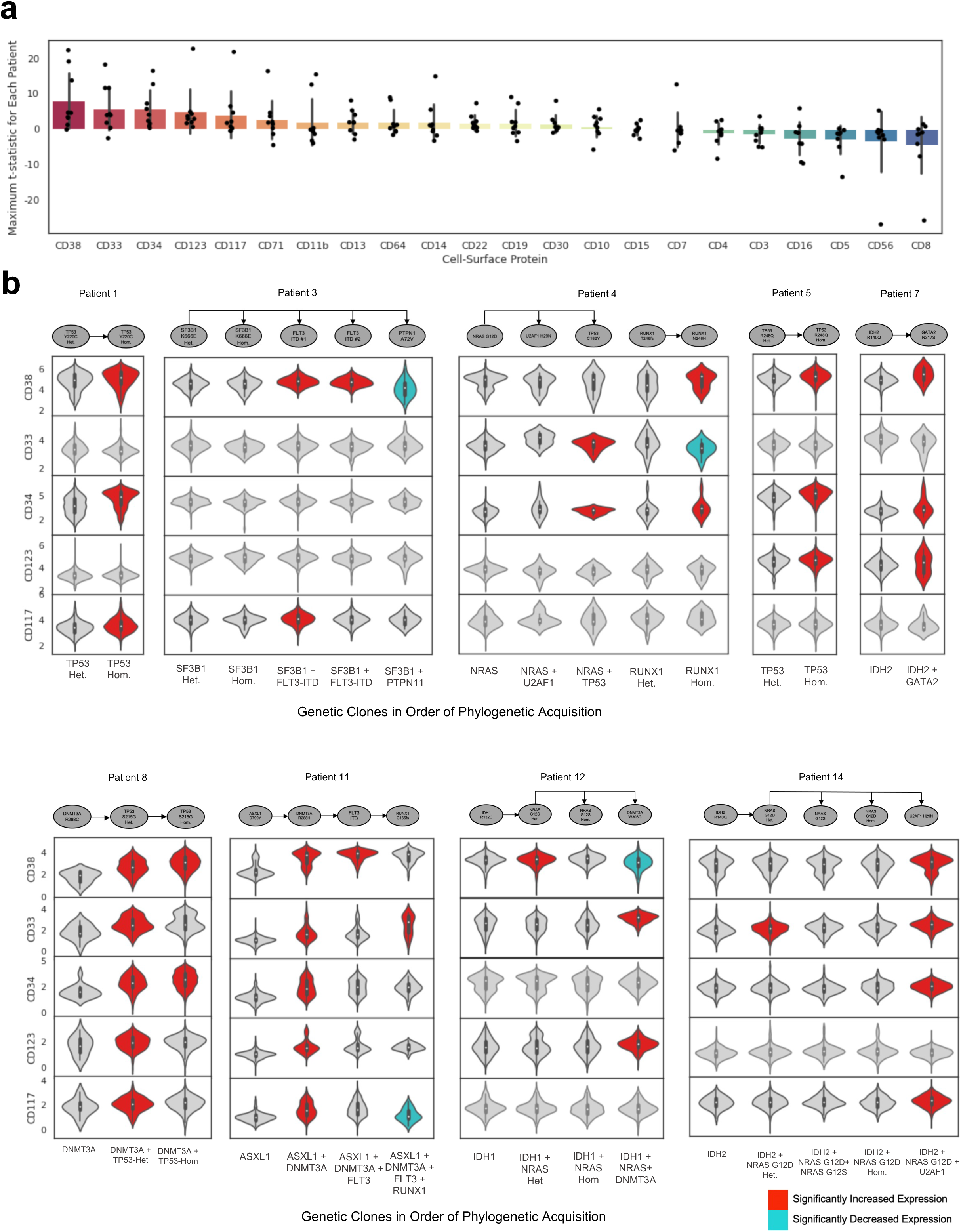
Association between immunophenotypic evolution and mutational acquisition. **A.** Box-and-whiskers plot with dot-plot overlain depicting maximum t-statistic for 22 cell-surface antibodies for each patient across all clones. For each antibody, antibody expression of all subsequent “branch” phylogenetic clones are compared to the founding phylogenetic clone, generating a t-statistic, and the maximum t-statistic for an individual antibody and patient is plotted. Each box represents 1 immunophenotypic protein and each overlain dot represents 1 of 9 individual patients. Immunophenotypic proteins are ranked by maximum t-statistic across all patients, ranging from CD38 (greatest increase in expression with mutational acquisition across patients) to CD8 (lowest increase in expression). **B.** *Top:* Mutation phylogeny of 9 patients with MPAL with at least 2 stepwise mutational acquisitions identified on single-cell DNA analysis. Each oval represents a genetically distinct subclone and arrows represent cumulative acquisition of mutational events. *Bottom:* Violin plots depicting expression of CD38, CD33, CD34, CD123, and CD117 for each subclone represented in the above phylogeny. Violin plots color-coded in red indicate protein expression that has significantly increased with mutational acquisition; plots color-coded in blue indicate a significant decrease in protein-expression. Statistical significance is considered p < 0.05 after adjustment via the Bonferroni method. Het: Heterozygous; Hom: Homozygous. All mutations are heterozygous unless specified otherwise.

Across all 9 patients, the maximal change in protein expression was greatest for CD38, CD34, CD33, CD123, and CD117, markers associated with immaturity (hematopoietic stem cells, and in some cases common myeloid or granulocyte-monocyte progenitor cells). Therefore, with progressive mutational acquisition, there was increased expression of these 5 markers of immaturity. Figure 3B depicts the change in expression of these 5 immunophenotypic proteins for all 9 patients. Despite containing diverse mutations, all 9 patients demonstrated significant increase in the expression of at least 2 of these 5 proteins with mutational acquisition, and in 2 patients (Patient 7 and Patient 11), expression of all 5 proteins increased. Furthermore, for patients with 3 or more stepwise mutational acquisitions, these immaturity markers often increased multiple times. For example, in Patient 4, CD34 expression significantly increased with acquisition of a *TP53* mutation, and then significantly increased again with subsequent acquisition of a *RUNX1* mutation. Collectively, these findings suggest that in MPAL leukemic progression, mutational evolution is associated with transition to a more immature immunophenotype.

While increased expression of immature markers CD38, CD34, CD33, CD123, and CD117 was the most common immunophenotypic change, evidence of cellular maturation and differentiation was seen in select genetic branches. For example, in Patient 12, acquisition of a terminal *DNMT3A* mutation was associated with increased expression of CD11b, CD13, CD14, and CD64, consistent with myeloid and monocytic differentiation (Supplementary Figure 7).

### Transcription and Immunophenotype Associate Incompletely Across Patients

We next examined how gene expression was associated with immunophenotype across all patients. Through unsupervised clustering of immunophenotypic markers across all cells and all patients, we identified 13 immunophenotypically-defined subpopulations. For many of these subpopulations, the cell type as identified by transcription closely associated with the expected immunophenotype (Supplementary Figure 8). For example, transcriptionally defined normal T cells were composed of 87.2% CD3+/CD5+ cells, while transcriptionally defined normal B cells were 94.2% CD19+/CD22+ cells (Figure 4A).

**Figure 4.**
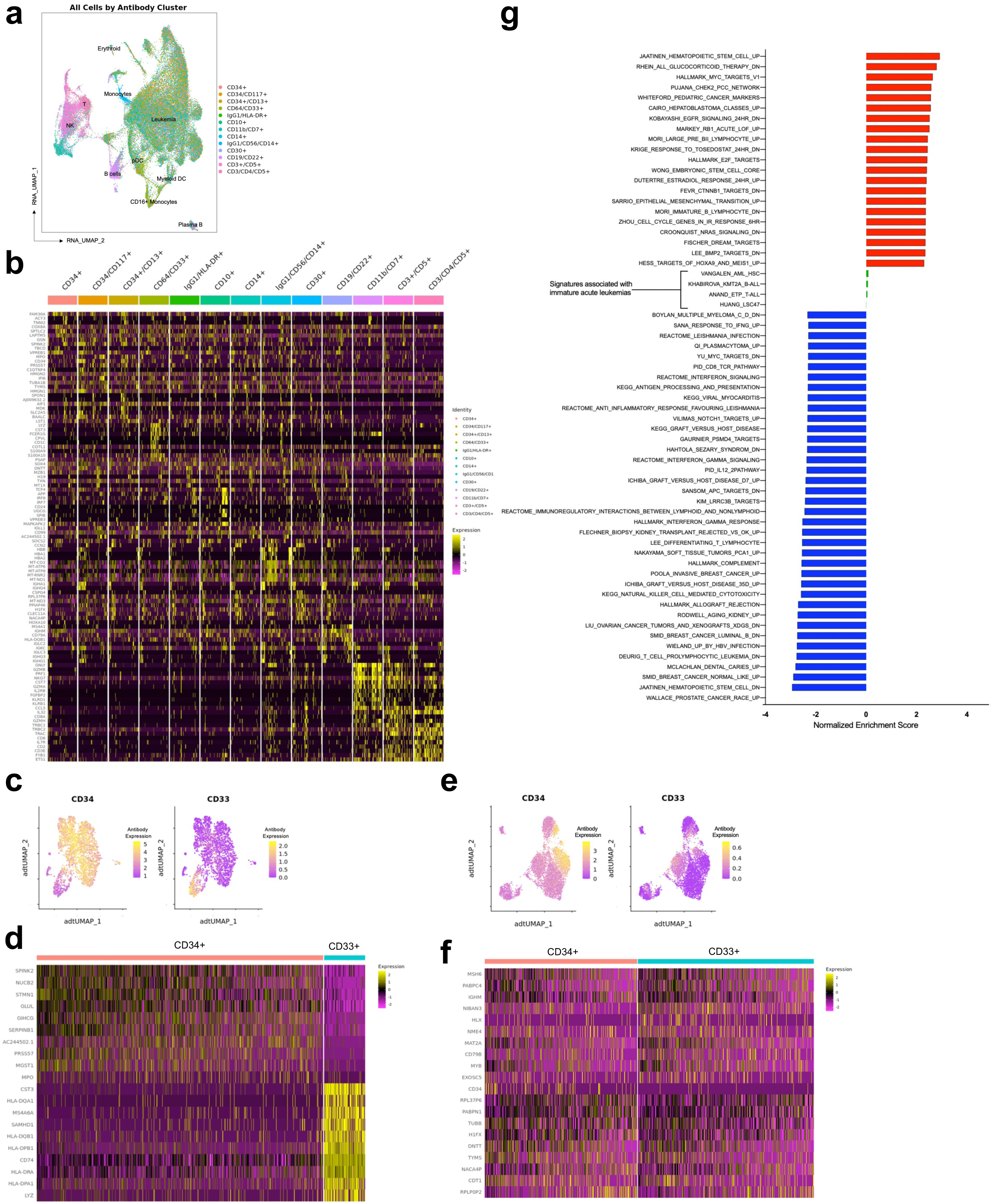
MPAL is comprised of heterogenous transcription-immunophenotypic associations and a common, stemlike transcriptional signature. **A.** RNA-derived UMAP from SC CITE-seq analysis of 71,579 cells from 12 patients with MPAL from Figure 1E. Cells are annotated based on transcriptionally defined cell populations. Cells were clustered by the expression of cell-surface immunophenotypic protein expression into 13 immunophenotype-defined clusters; cells were then color-coded based on cluster. **B.** Heatmap of scaled expression values for top ten most upregulated genes in each of the 13 immunophenotypic subpopulations from 4A. **C.** RNA-derived UMAP from 2,594 cells from **Patient 11**. Cells are color-coded based on expression of CD34 (*left*) and CD33 (*right*). **D.** Heatmap of scaled expression values for top ten most upregulated genes for the CD34- positive cell population (*left columns*) and the CD33-positive cell population (*right columns*) from Patient 11. **E.** RNA-derived UMAP from 6,100 cells from **Patient 2**. Cells are color-coded based on expression of CD34 (*left*) and CD33 (*right*). **F.** Heatmap of scaled expression values for top ten most upregulated genes for the CD34- positive cell population (*left columns*) and the CD33-positive cell population (*right columns*) from Patient 2. **G.** Bar plot of normalized enrichment scores of 59 gene sets with false-discovery rate q-values < 0.00005 identified via gene set enrichment analysis (GSEA) of all cells transcriptionally annotated as leukemia vs non-leukemia across 12 patients with MPAL. Positively enriched gene sets are color-coded in red and negatively enriched gene sets are color coded in blue. As a comparison, 4 gene sets previously described in immature AML or ALL are also shown and color-coded in green; these genesets are not enriched.

To contrast, the transcriptionally defined ‘leukemia’ cells were comprised of cells from heterogeneous immunophenotypic subpopulations, with the greatest contributions from cells with stem or myeloid markers, including CD34+/CD13+ cells (12.89% of leukemia population), CD34+/CD117+ cells (12.86%), IgG1+/HLA-DR+ cells (11.81%), CD33+/CD64+ cells (11.60%), and CD34+/CD33+/CD117+ cells (11.20%). Cells with lymphoid markers were also present in the transcriptionally defined leukemia cells, but in smaller proportions with CD19+/CD22+/CD30+ cells comprising 5.96% of the leukemia cluster, followed by CD19+/CD22+/CD45+ cells (5.49%), CD3+/CD5+/CD7+ cells (4.45%), and CD3+/CD4+/CD5+ cells (0.4%) (Figure 4A). While the immunophenotypic subpopulations demonstrated some differences in gene expression, many had similar expression patterns, reflecting that many individual cells had similar gene expression despite having heterogeneous immunophenotypes (Figure 4B).

### Transcription and Immunophenotype Associate Incompletely Within Individual Patients

On the individual patient level, the association between transcription and immunophenotype was heterogeneous. In some patients, immunophenotype was closely associated with a distinct transcriptional signature. For example, in Patient 11, immunophenotype-based clustering revealed distinct CD34+ and CD33+ populations (Figure 4C). In addition to having distinct immunophenotypes, these two populations also had distinct gene expression profiles, with the CD33+ population demonstrating markedly higher expression of major histocompatibility complex-encoding genes *HLA-DQA1, HLA-DQB1, HLADPB1, HLA-DRA*, and *HLA-DPA1,* relative to the CD34+ population (Figure 4D). In other patients, however, immunophenotype and transcriptional profile were not closely associated. For example, in Patient 2, immunophenotype-based clustering also revealed distinct CD34+ and CD33+ subpopulations (Figure 4E). Unlike Patient 11, however, in Patient 2 these two immunophenotypically-distinct subpopulations did not have distinct gene expression profiles (Figure 4F).

### MPAL cells upregulate stemlike pathways and are distinct from other acute leukemias

To further define the transcriptional signature of MPAL, we performed gene set enrichment analysis (GSEA) on all transcriptionally annotated as leukemia cells across all patients using the molecular signature database (mSigDB) hallmark and C2 gene sets (Figure 4G)^11, 12^. GSEA demonstrated enrichment for gene sets associated with stem cells. The greatest enrichment was demonstrated for a gene signature first described in CD133+ stem cells derived from human cord blood (normalized enrichment score [NES] 2.92, q-value 0.0); genes associated with embryonic stem cells were also highly enriched (NES 2.41) (Supplementary Figure 9A)^13, 14^. Decreased enrichment was demonstrated in gene signatures associated with immune or inflammatory pathways, including natural killer cell cytotoxicity, complement activation, and interferon gamma signaling (Supplementary Figure 9B).

In addition to mSigDB gene sets, we also assessed enrichment of gene sets derived from transcriptional analysis of other immature leukemias. For this analysis, we included gene signatures associated with early T-cell progenitor (ETP) ALL, *KMT2A-*rearranged B-cell ALL, hematopoietic-stem cell (HSC)-like AML, and the leukemia stem cell (LSC)-47, a gene score associated with immature AML^15–18^. We also compared gene signatures derived from more differentiated acute leukemias, including granulocyte-monocyte progenitor-like AML, myeloid-like AML, and *NUTM1-*rearranged ALL^15, 17^. None of these AML or ALL-derived gene signatures were enriched in MPAL above a threshold of NES> +/- 0.1 (nominal p-value > 0.05) (Figure 4G), indicating that the MPAL transcriptional profile is distinct from known signatures associated with either lymphoid or myeloid acute leukemias.

### MPAL cells demonstrate variable differentiation potential which predicts survival

Given enrichment for genes associated with stemness as well as the lack of enrichment of known leukemia gene signatures, we sought to apply a more recently developed metric of stemness, CytoTRACE [for cellular (Cyto) Trajectory Reconstruction Analysis using gene Counts and Expression]^19^, to our SC transcriptional dataset. CytoTRACE is a computational framework for predicting the differentiation potential of a single cell based on transcriptional data about numbers of expressed genes, covariant gene expression, and local neighborhoods of transcriptionally similar cells. CytoTRACE provides a score for each cell representing its stemness within a given dataset,^19^ ranging from 0 to 1, with higher scores indicating greater stemness. When applied to our 12 patient cohort, we found high CytoTRACE scores to be overrepresented in our ‘leukemia’ cluster relative to non-leukemic populations (median CytoTRACE 0.61 vs 0.23 for leukemia vs non-leukemia populations, p < 2e-16) (Figure 5A).

**Figure 5.**
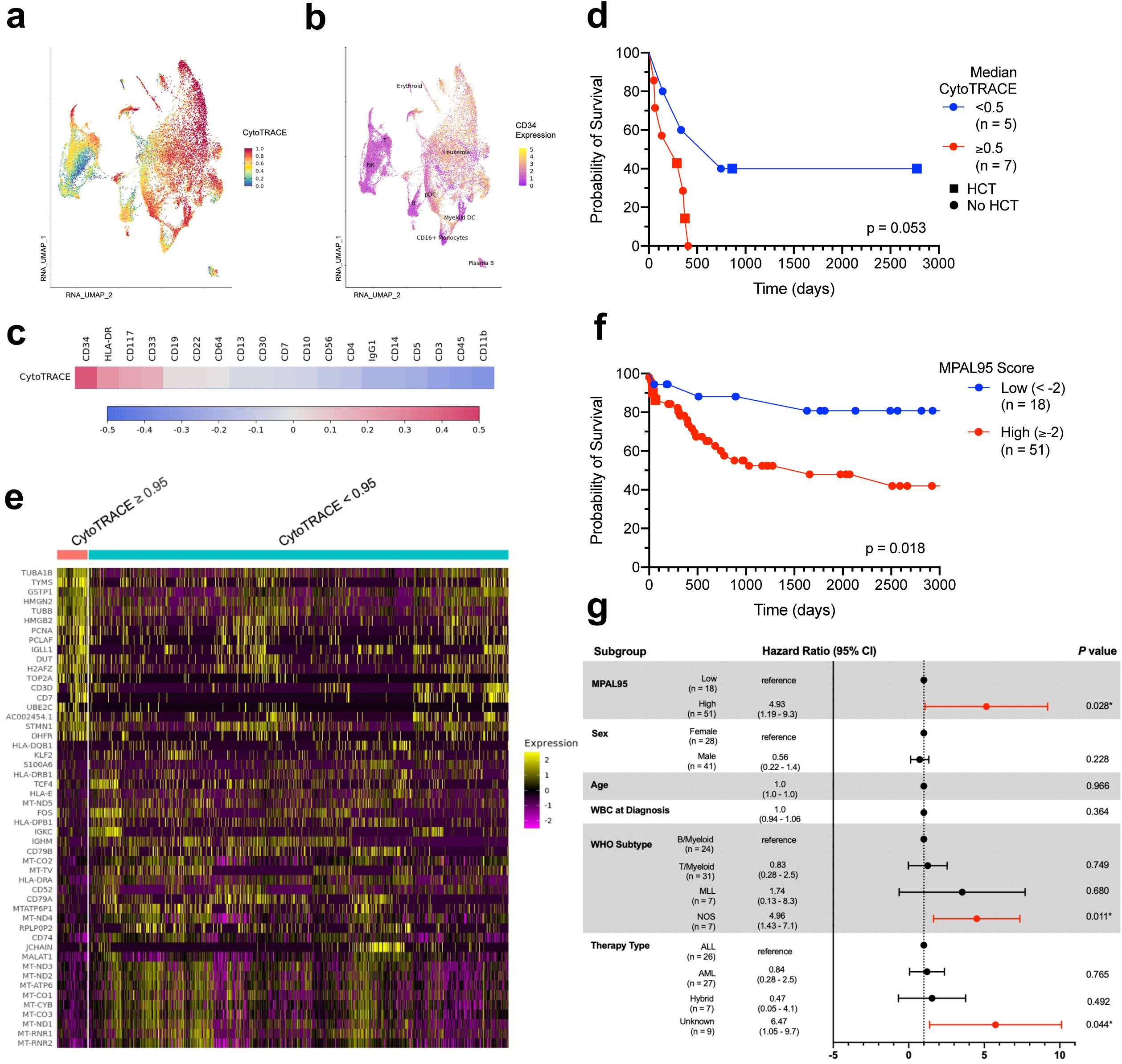
**MPAL cells are stemlike and measures of stemness are prognostic of patient outcomes** **A.** RNA-derived UMAP from SC CITE-seq analysis of 71,579 cells from 12 patients with MPAL from Figure 1E. Cells are color-coded based on cytoTRACE score from 0 (most differentiated) to 1 (least differentiated). **B.** UMAP from 5A. Cells are color-coded based on cell-surface expression of CD34 protein. **C.** Spearman correlation matrix of CytoTRACE score and cell-surface protein expression. Correlation coefficient is denoted by color coding. **D.** Kaplan-Meier estimates of overall survival stratified by median CytoTRACE score <0.5 vs >0.5 for 12 adult patients with MPAL. **E.** Heatmap of scaled expression values for the genes with greatest upregulation in single cells with high cytoTRACE (> 0.95) (*left columns*) vs low cytoTRACE (<0.95) (*right columns*). **F.** Kaplan-Meier estimates of overall survival stratified by MPAL95, a gene set score derived from single-cell transcriptional data, for 69 pediatric patients with MPAL from the TARGET initiative. **G.** Multivariate Cox proportional hazards model for 69 pediatric patients with MPAL, with the MPAL95 gene signature included. CI, confidence interval.

Across the cohort, CytoTRACE score was moderately correlated with higher CD34 expression, followed by HLA-DR, CD117, and CD33 expression (Spearman correlation coefficient 0.44, 0.25, 0.20, 0.18 for CD34, HLA-DR, CD117, and CD33, respectively) (Figure 5B-C). For individual patients, the median CytoTRACE score of each patient’s leukemia population varied considerably, ranging from 0.13 (least stemlike) to 0.89 (most stemlike). When stratified by median CytoTRACE score of the leukemia population, a higher median CytoTRACE trends towards an inferior overall survival (OS) in our small cohort (p = 0.053) (Figure 5D).

Relative to single cells with lower CytoTRACE scores (< 0.95), single cells with very high CytoTRACE scores (>= 0.95) demonstrated a distinct gene expression profile (Figure 5E). In a GSEA, cells with CytoTRACE scores >= 0.95 demonstrated upregulation of multiple pathways associated with cellular proliferation, cell cycle dysregulation, and a stem or progenitor-like cell state (Supplementary Figure 10).

### Generation of a CytoTRACE-based prognostic signature and validation in an independent cohort

We next sought to derive and validate a CytoTRACE-based prognostic metric in an independent cohort of patients with MPAL. To generate a CytoTRACE-based score, we compared the differential gene expression of single cells with very high (>= 0.95) vs low (<0.95) CytoTRACE scores. Genes with greatest upregulation in the cells with high CytoTRACE scores were then used to compute a gene set score, which we termed MPAL95. When pseudo-bulking was applied to the leukemic populations of our cohort, we confirmed that MPAL95 was prognostic for OS while the LSC-17, a transcriptionally-based risk stratification system previously described in AML^18^, was not (Supplementary Figures 11A-B).

The prognostic ability of MPAL95 was validated using external bulk RNAseq data of acute leukemias of ambiguous lineage from the Therapeutically Applicable Research To Generate Effective Treatments (TARGET) initiative, which includes expression profiles for 115 pediatric patients with MPAL; 69 patients with available survival data were included in this analysis^20^. Patients from the TARGET cohort demonstrated variable MPAL95 scores (Supplementary Figure 11C). Relative to patients with the lowest MPAL95 scores, patients with high MPAL95 scores demonstrated significantly inferior OS with a 2-year OS of 62.6% (95%CI 50.2% - 78.1%) for patients with high MPAL95 scores versus 88.1% (95% CI 73.9% - 99.9%) for patients with low MPAL95 scores (p = 0.018; Figure 5E, Supplementary Figure 11D). This relationship was preserved in a multivariable Cox regression model. High MPAL95 score was significantly associated with inferior OS independent of patient age, sex, white blood cell count at diagnosis, WHO subtype, and type of frontline treatment, with a hazard ratio of 4.93 (95% confidence interval 1.19–- 20.3, p = 0.028) (Figure 5F). By contrast, the LSC-17 was not prognostic for OS in the TARGET cohort (Supplementary Figure 11E).

## Discussion

There is a critical need for improved outcomes in MPAL. The historical lack of biologic understanding and subsequent confusion in defining this disease entity remain critical barriers to improving survival. There are no consensus guidelines for treatment. In current practice, patients are treated with either ALL- or AML-like chemotherapy, a decision often made arbitrarily rather than driven by disease biology. A recent analysis suggested matching treatment to ALL- or AML-like chemotherapy based on methylation profiles may improve remission rates, but this has not been adopted into clinical practice^5^. Without appropriate definition and sub-classification of MPAL, clinical trials to optimize therapy are challenging. Furthermore, no risk stratification for MPAL currently exists. In this context, we offer a dissection of the cellular genetic and transcriptional origin of MPAL and provide a framework for disease definition and risk stratification. Our SC multiomic analysis of newly diagnosed MPAL samples allows for direct measurement of cell surface markers comprising the ‘mixed’ immunophenotype and permits explicit correspondence of immunophenotype with genetic and transcriptomic profiles.

Characterization of these relationships at the SC level has not previously been performed in MPAL. Our data contradicts the assumption that MPAL lies on a continuum with other acute leukemias, and instead classifies MPAL as a distinct leukemia with stem-like features. Unlike ALL and AML, which are defined and risk stratified by genetics, we distinguish MPAL with a transcription-based metric that correlates with patient survival and can be distilled into a score, MPAL95, which permits risk stratification from bulk RNAseq data.

Our data provides novel insight into the biology and leukemogenesis of MPAL. The gene mutation panel in our study was not exhaustive, including a limited set of 20 genes commonly mutated in acute leukemias. Nonetheless, we observed stepwise acquisition of mutations in 9 patients and associate mutation phylogeny with immunophenotypic changes. Most leukemias are thought to be driven by a series of successive genetic alterations, culminating in transformation to malignant disease. This canonical road of leukemogenesis, when applied to MPAL, suggests that sequential mutation acquisition leads an MPAL cell to have increased potential for lineage plasticity. Prior investigation into MPAL biology suggested the stem-like nature of MPAL and proposed that mutation in a multipotent progenitor cell leads to lineage promiscuity^3^. Our data also supports the stem-like nature of MPAL. As defined by CytoTRACE, MPAL cells express a stem-like gene expression profile and demonstrate a large number of expressed genes.

We have shown that, in MPAL, markers of immaturity can be gained alongside mutations over time. In our data, mutational acquisition was associated with increased expression of multiple cell-surface proteins associated with an immature and less differentiated cell state. MPAL may, therefore, arise from a primitive cell, or an MPAL cell may revert to a more primitive phenotype with successive mutational evolution. This suggests that the MPAL cell of origin may span a spectrum of differentiation and supports that a cell’s leukemic potential cannot be assigned by immunophenotype. Microenvironmental factors or epigenetics may also influence the translation of the genome or transcriptome to lineage expression in individual leukemic populations (Supplementary Figure 12).

This improved biologic understanding further demarcates MPAL as distinct from ALL or AML. MPAL shares many common genetic aberrations with AML and ALL. In our MPAL dataset, as in SC AML data, signaling mutations were not co-mutated within the same cell or clone, suggesting that some co-mutational patterns may be conserved across histologies ^7, 8, 21^. The phenotypes resulting from specific mutations, however, may be discrete to each leukemia. For example, across our MPAL cohort, *NRAS* mutations were associated with a more immature phenotype with increased CD34 and CD38 cell-surface protein expression. This is in contrast with *NRAS* mutations in AML, which are associated with a more monocytic immunophenotype^22, 23^. Further, the gene expression signature that distinguishes MPAL is distinct from those previously described in ALL or AML or ALL, including stem-like AML and ETP-ALL, considered the most primitive pure lymphoid leukemia^15–18^. The distinct biology of MPAL likely means that it will require its own unique sub-classification, which cannot be extrapolated from other leukemias.

Importantly, we define a novel prognostic gene signature for MPAL. Although the nomenclature of MPAL suggests that the ‘mixed phenotype’ is the most salient disease component, our data challenges this assumption. Our data suggest that the mixed immunophenotype of MPAL, while demonstrative of lineage derangement, may have little clinical relevance. Instead, the transcriptional signature and the degree of differentiation potential represented by this signature likely determine clinical behavior. More specifically, differentiation potential as measured by CytoTRACE correlates with a more proliferative, aggressive leukemia, and predicts survival. From SC data, we then derived MPAL95, a prognostic gene set score applicable to bulk RNAseq that predicts survival in an independent validation cohort.

This work lays the foundation for a MPAL-specific risk stratification system, which does not currently exist, and supports prospective validation of transcriptionally defined stemness as a prognostic biomarker. Since our study is retrospective and MPAL is a rare disease, our sample size is small. Despite this, we have the largest cohort of adult MPAL patients assessed by SC analysis to date. Validation of our derived bulk prognostic score in an independent pediatric cohort^20^ spanning multiple decades and treatment strategies supports the validity of our model of MPAL as a disease both defined and prognosticated by differentiation potential. Further, the fact that our prognostic score holds in pediatric and adult patients emphasizes the importance of stemness over other disease factors, including genetics, treatment approach, and age^3, 5^. Future clinical studies will be needed to validate CytoTRACE and MPAL95 as prognostic tools and to elucidate optimal treatment strategies for MPAL across the span of differentiation potential. Finally, further mechanistic studies will be required to characterize the true cell of origin for MPAL and determine the genetic, epigenetic and microenvironment factors that drive stemness and disease behavior.

## Methods

### Patient Samples

Cryopreserved bone marrow or peripheral blood mononuclear cells from 14 adult patients with newly diagnosed MPAL were included in this study. Patients were diagnosed at either the University of California San Francisco or the University of Pennsylvania from 2006-2020, and initial diagnosis was made pathologically using World Health Organization criteria operative at the time of diagnosis. All patients provided written informed consent for sample banking and analysis under protocols approved by the local Institutional Review Board and conducted in accordance with the ethical standard of the institution and with the Declaration of Helsinki. All 14 samples were analyzed with simultaneous SC DNA and cell surface protein sequencing and 12 samples were concurrently analyzed with SC RNA and cell surface protein sequencing (Figure 1A).

### Single-cell DNA and Protein Sample Preparation, Library Generation, and Sequencing

We performed DAb-seq on unsorted mononuclear cells from 14 patients using a microfluidic approach with molecular barcode technology using the Tapestri platform (MissionBio) as previously described^6, 24^. Briefly, cryopreserved cells were thawed, normalized to 10,000 cells/μL in 180 μL PBS (Corning), and then incubated with Human TruStain FcX (BioLegend) and salmon sperm DNA (Invitrogen) for 15 minutes at 4C. A pool of 22 oligo-conjugated antibodies (CD3, CD4, CD5, CD7, CD8, CD10, CD11b, CD13, CD14, CD15, CD16, CD19, CD22, CD30, CD33, CD34, CD38, CD56, CD64, CD71, CD117, CD123) (Supplementary Table 3) was added, and cells were incubated for an additional 30 minutes. In addition, individual samples were also incubated with unique anti-CD45 oligo-conjugated antibodies to provide sample-level identifiers, and groups of 3 individual patients were pooled together for multiplexed runs. All oligo-conjugated antibodies were generated as previously described and were run on a Bioanalyzer Protein 230 electrophoresis chip (Agilent Technologies, cat. no 5067-1517) to verify successful conjugation^6^.

Next, pooled samples were resuspended in cell buffer (MissionBio), diluted to 4-7e6 cells/mL, and then loaded onto a microfluidics cartridge, where individual cells were encapsulated, lysed, and barcoded using the Tapestri instrument. DNA from barcoded cells was amplified via PCR using a targeted panel that included 127 amplicons across 20 genes associated with acute leukemia (Supplementary Table 2). DNA PCR products were isolated, purified with AmpureXP beads (Beckman Coulter), used as a PCR template for library generation, and then repurified with AmpureXP beads. Protein PCR products were isolated from the supernatant from AmpureXP bead purification via incubation with a 5’ Biotin Oligo (ITD). Protein PCR products were then purified using Streptavidin C1 beads (Thermo Fisher Scientific), used as a PCR template for library generation, and then repurified using AmpureXP beads. Both DNA and protein libraries were quantified and assessed for quality via a Qubit fluorometer (Life Technologies) and Bioanalyzer (Agilent Technologies) prior to pooling for sequencing on an Illumina Novaseq.

### Single-Cell DAb-seq Data Processing and Analysis

FASTQ files from single-cell DAb-seq samples were processed via an open-source pipeline as described previously^6, 25^. This analysis pipeline trims adaptor sequences, demultiplexes DNA panel amplicons and antibody tags into single cells, and aligns panel reads to the hg19 reference genome. Valid cell barcodes were called using the inflection point of the cell-rank plot in addition to the requirement that 60% of DNA intervals were covered by at least eight reads. Variants were called using GATK (v 4.1.3.0) according to GATK best practices^26^. ITDseek was used to detect *FLT3* internal tandem duplication (ITD) from amplicon sequencing reads^27^. For each valid cell barcode, variants were filtered according to quality and sequence depth reported by GATK, with low quality variants and cells excluded based on the cutoffs of quality score < 30, read depth < 10, and alternate allele frequency < 20%. Cell-surface protein reads were normalized using centered log ratio transformation^28^.

### SNP and Antibody-Based Demultiplexing

To de-multiplex individual patients combined into a single sample, we used a novel computational approach incorporating both patient-specific oligo-conjugated “hash” antibody tags as well as single nucleotide polymorphisms (SNPs) covered by the SC DNA panel^29^.

Individual patient samples were stained with unique oligo-conjugated “hash” antibodies and then multiplexed into groups of 3. All SNPs were treated as binary (mutated or wildtype). To identify SNPs that maximally differ between samples, for each multiplexed group, we filtered all SNPs mutated in <10% or >80% of cells. For the remaining SNPs, missing data was imputed based on a majority vote of the binary data from the 5 nearest neighbors using the kNN function from the VIM package in R. Next, we hierarchically cluster cells using cosine as the distance function and Ward’s method for joining clusters and cut the resulting dendogram into 3 clusters, one for each patient. To refine the SNPs included in clustering, Fisher’s exact test was computed between the SNP value and cluster membership across cells; SNPs with p-values < 10^-12^ were selected and re-clustered in the same hierarchical manner.

Next, SNP-based cell clusters were refined using hash antibody data. Starting with 3 SNP-based clusters, we add additional clusters by traversing down the hierarchical tree and splitting if there was a significant difference between the current cluster and subsequent split by Hotelling’s T2 test with a p-value cutoff of 10^-5^. Splitting was stopped when there were <10 cells per cluster. Clusters were then assigned to a specific hash antibody by comparing the antibody expression of the cluster to the expected hash background distribution. For each hash antibody, the antibody expression for a multiplexed experiment is expected to be bimodal, with one right mode comprised of antibody-stained cells belonging to a single patient and one left mode comprised of unstained cells. To estimate the expected background antibody distribution, we generated a symmetric distribution by reflecting the data to the left of the left mode about the mode. Clusters were assigned to a specific hash antibody and patient if >50% of cells from that cluster demonstrated hash antibody expression above the 95^th^ percentile of the expected background distribution. A cluster was considered a multiplet if it was assigned to multiple patients. Cells designated as multiplets or unassignable were excluded from downstream analyses.

### Clonal Analysis and Inference of Mutational Phylogenies

Following de-multiplexing, for individual patients, we analyzed all variants present in >0.1% of cells. Variants were assessed for known or likely pathogenicity via ClinVar and COSMIC databases^30, 31^, and previously identified, nonintronic somatic variants were included in clonal analyses. Genetic clones were defined as >10 cells possessing identical genotype calls for the protein encoding variants of interest, as per prior SC DNA studies^7, 21^. Phylogenetic trees for individual patients were inferred using single cell inference of tumor evolution (SCITE), a probabilistic model for inferring phylogenetic trees using a flexible Markov-chain Monte Carlo algorithm^10^. SCITE was employed with a global false positive rate set to 1% and a platform-provided false-negative rate, as per prior SC DNA studies^8^. To define immunophenotypic subpopulations, unsupervised hierarchical clustering was performed using the *scipy* package in python on scaled and centered log ratio-normalized protein expression data. UMAPs derived from protein expression data were constructed using the *umap* function in python with default settings.

We further measured how cell-surface immunophenotypic protein expression changed with progressive acquisition of mutations in the 9 patients of our 14 patient cohort that had at least 2 stepwise mutational acquisitions identified on SC phylogenetic analysis (Supplementary Figure 1). To do this, for each patient, we compared expression of each of the 22 immunophenotypic proteins for the founding genetic clone to all subsequent genetic clones and calculated a t-statistic. To identify which cell-surface proteins changed the most with mutational acquisition across the cohort as a whole, for each patient, we determined the maximum t-statistic for each immunophenotypic protein (Figure 3A).

### Single-cell RNA and Protein Sample Preparation, Library Generation, and Sequencing

We performed SC CITE-seq sequencing on unsorted mononuclear cells from 12 patients using a particle-templated instant partitions sequencing (PIPseq) platform^9^. Briefly, cryopreserved cells were thawed, and 1-2 million cells were incubated in 45ul of Cell Staining Buffer (BioLegend) per million cells with Trustain FcX block (BioLegend) for 15 minutes on ice. A pool of 19 oligo-conjugated antibodies (CD3, CD4, CD5, CD7, CD10, CD11b, CD13, CD14, CD19, CD22, CD30, CD33, CD34, CD45, CD56, CD64, CD117, IgG1, HLA-DR) were added and incubated on ice for an additional 60 minutes. Cells were quantified, resuspended in PBS with 0.04% BSA, and combined in a 1:10 ratio with barcoded hydrogel templates (1000 cells/ul) and processed according to PIPseq Single Cell Epitope Sequencing Use Guide Rev 2.0 (FB0002079). Briefly, Partitioning Reagent (Fluent BioSciences) was added to the cell-PIP mixture and vortexed on a custom vortexer (Fluent BioSciences). After removal of excess Partitioning Reagent, the emulsion was placed on a dry bath (66°C for 40 minutes followed by 4°C for 11 minutes) for cell lysis and RNA capture. Emulsions were broken with De-Partitioning Reagent (Fluent Biosciences), washed, and cDNA synthesis was conducted on the RNA hybridized to PIP templates in bulk. Double-stranded DNA libraries were then enzymatically fragmented and adapters for Illumina sequencing were ligated prior to amplification with appropriate index adapters. The resulting PIPseq libraries were pooled and sequenced using an Illumina NextSeq2000.

### Single-Cell CITE-seq Data Processing and Analysis

FASTQ files from single-cell CITE-seq were processed via PIPseeker v0.52 (Fluent). This RNA pipeline comprises 4 basic steps: barcode identification and error correction, mapping to the reference transcriptome, gene expression matrix generation, and cell calling. Adapter sequences are trimmed, data is demultiplexed into single cells (BCL Convert, Illumina Basespace dashboard), matched against a list of known barcodes and mapped (Salmon alevin v1.4.0) against the GRCh38.p13 reference transcriptome, and putative cells are separated from background^9^. Genes were filtered if detected in <3 cells and cells were filtered based on having low-complexity libraries (feature count < 200) or high mitochondrial content (>15%). Similarly, ADT analysis was also processed via PIPseeker v0.52 (Fluent), including error correction, trimming of adapter sequences, mapping to a list of known barcodes, and generating a UMI matrix (CITE-seq Count v1.4.3). Downstream bioinformatics analysis, including scaling and log-normalization of single-cell transcriptional data and centered-log ratio normalization of single-cell protein expression data, was performed using Seurat 4.3.0^32^. Across all samples, data integration was performed using reciprocal principal component analysis (RPCA). Unsupervised cell clustering on transcriptional data was performed using Seurat with resolution set to 0.6, and clusters were visualized using the Seurat function *RunUMAP* with default settings. Cell populations were annotated by RNA expression using a combination of scType and clustifyr followed by independent manual confirmation via marker genes^33, 34^. Both annotation frameworks agreed on all clusters apart from a population of cells assigned as “cancer cells”, “pro-B cells”, “progenitor cells”, or “unknown” by scType and “CD34+” cells by clustifyr; this cluster was collapsed into a common “leukemia” cluster. Differentially expressed genes for each cluster were determined using Seurat’s *FindConservedMarkers, FindAllMarkers* or *FindMarkers* functions, as appropriate.

### Gene Set Enrichment Analyses

Gene set enrichment analyses (GSEA) were performed using gsea v4.2.3 on genes pre-ranked by log2 fold change generated by comparing cells annotated as leukemia vs non-leukemia or by comparing leukemia cells with CytoTRACE ≥ 0.95 vs < 0.95^35^. Gene sets used in this study included the molecular signatures database (MSigDB) hallmark v2022.1 (50 genes) and the c2 gene set curated from various sources in the biomedical literature (6,449 genes)^11, 12^. To compare MPAL to other leukemias, in the GSEA comparing cells annotated as leukemia vs non-leukemia, we also included the following 6 gene signatures associated with: early T-cell progenitor (ETP) ALL, *KMT2A-*rearranged B-cell ALL, *NUTM1-*rearranged ALL, hematopoietic-stem cell (HSC)-like AML, granulocyte-monocyte progenitor-like AML, and the leukemia stem cell (LSC)-47, a gene score associated with immature AML^15–18^.

### CytoTRACE-Based Analyses

Differentiation potential was determined using CytoTRACE v0.3.3, with 3,000 single cells sub- sampled from the 12 individual patients^19^. To generate MPAL95, a CytoTRACE-derived gene set score, we compared the differential gene expression of single cells with a high CytoTRACE score (≥ 0.95) vs a low CytoTRACE score (<0.95). Genes with greatest upregulation in the cells with high cytoTRACE scores were used to compute a gene set score, called MPAL95, using the first principal component, in an approach similar to that used to compute gene set scores from single-cell transcriptional data in acute myeloid leukemia^23^. MPAL95 was then applied to bulk RNAseq data for 69 patients from the TARGET-ALL-P3 dataset; samples were only included if survival outcomes were available^20^. Additional clinical variables pulled from the TARGET dataset and included in multivariable survival analysis were patient age, sex, white blood cell count at diagnosis, disease classification per WHO classification, and treatment type, classified per TARGET as AML-like, ALL-like, hybrid, or unknown. As additional validation, MPAL95 was applied to pseudo-bulked RNAseq data derived from SC RNAseq data from the 12 adult patients in our cohort. To pseudo-bulk our data, we first sub-setted the transcriptionally-identified leukemic cell populations, extracted raw counts after quality filtering, and then aggregated counts to the sample level.

### Statistics and Reproducibility

Continuous variables were compared using Student’s *t*-test or Mann-Whitney U tests and categorical variables were compared using chi-squared or Fisher’s exact tests. To evaluate clone-level cooccurrence, a contingency table was constructed for each mutation pair and the log2-transformed odds ratio computed; Fisher’s exact test was used to evaluate statistical significance. The association between individual mutations and cell-surface antibody expression was determined using point-biserial correlations and the association between CytoTRACE and cell-surface antibody expression was determined using Spearman’s correlation. Survival analysis was estimated using Kaplan-Meier curves and compared using log-rank tests. Hazard ratios were calculated using the multivariable Cox proportional hazards model. All p-values for single-cell level comparisons were adjusted via the Bonferroni methods unless otherwise specified. All statistical analyses were performed in R (v. 4.0.2).

### Data and Code Availability

The data discussed in this publication have been deposited in NCBI’s Gene Expression Omnibus^36^ and are accessible through GEO series accession number GSE232074 (https://www.ncbi.nlm.nih.gov/geo/query/acc.cgi?acc=GSE232074). All downstream analysis scripts and processed data files are available at github.com/SmithLabUCSF/MPAL.

## Acknowledgements

Sequencing was performed at the UCSF CAT, supported by UCSF PBBR, RRP IMIA, and NIH 1S10OD028511-01 grants. This research was supported in part by the West Charitable Trust. CACP is supported (in part) by the National Cancer Institute of the National Institutes of Health under Award Number K12CA260225. The content is solely the responsibility of the authors and does not necessarily represent the official views of the National Institutes of Health. CCS is the Damon Runyon-Richard Lumsden Foundation Clinical Investigator supported (in part) by the Damon Runyon Cancer Research Foundation (CI-99-18).

