## Supplemental Files for "Multiomic Single Cell Sequencing Identifies Stemlike Nature of Mixed Phenotype Acute Leukemia and Provides Novel Risk Stratification"

### Supplementary Table 1: Patient Demographics, Disease Characteristics, and Clinical Outcomes

| Patient Number | Age at MPAL Diagnosis | Sex | Clinical Subtype | Clinical Flow Cytometry | Cytogenetics | Clinical Bulk Next Generation Sequencing | Frontline Therapy | Clinical response to frontline therapy | Allogeneic HCT? | Survival After MPAL Diagnosis (days) | Alive? |
|---|---|---|---|---|---|---|---|---|---|---|---|
| 1 | 74 | M | B/Myeloid | Expressed CD7, CD13, CD34, CD38, variable CD56, weak CD71, CD117, HLA-DR, Subset co-expressed MPO (9% of gated events positive), Subset co-expressed CD19 (10% of gated events positive), Subset co-expressed TdT (59% of gated events positive), Subset co-expressed cytoplasmic CD79a (46% of gated events positive). | Complex: 45-46,XY,add(5)(q22),add(6)(q13)[5]/44,sl,-7,add(16)(q22),add(17)(p11.2),add(19)(q13.3),add(20)(q11.2),-21[3]/44,sdl1,+der(19)t(1;19)(q21;q13.4)dup(1)(q21q32), der(21)t(11;21)(q13;p11.2),-22[3]/44,sdl1,+?add(19)(q13.1),-22[7]/86-88,sdl2x2[2]. | TP53 T220C (VAF 67%), CSF3R S810fs (VAF 13%), CSF3R Q768% (VAF 3%) | Decitabine | Refractory disease | N | 143 | N |
| 2 | 61 | F | B/Myeloid | Blasts expressed CD15, CD19, CD22, cytoplasmic CD79a, HLA-DR, TdT; subset also expressed CD11c (9%), subset expressed CD34 (26%); subset expressed CD64 (15%); rare events expressed CD34 and CD117, consistent with myeloid blasts. | Normal female karyotype | Not available | Daunorubicin, vincristine, prednisone, peg-asparaginase | CR | Y | 2774 | Y |
| 3 | 78 | M | B/Myeloid | Blasts Expressed: CD34-/ CD19-/ CD10-/ CD20 var/ HLADR bright /(n)TdT+/ (c)CD22+/ (c)CD79a+. This population also shows some aberrant dim expression of CD4 and CD2, along with a subset of the region positive for CD7 | 46,XY,+1,der(1;14)(q10;q10 | DNMT3A R736H (VAF 48%), FLT3-ITD c.1781(VAF 32%), FLT3-ITD c.1813 (VAF 7%), SF3B1 K666E (VAF 50%), TET2 H864Q (VAF 52%) | Sorafenib, methotrexate, mercaptopurine | CRi | N | 405 | N |
| 4 | 49 | M | B/Myeloid | Expressed weak-variable CD15, CD19, weak CD22, variable CD34, CD38, weak CD71, weak CD123, variable HLA-DR, Subset co-expressed CD56 (24% of gated events positive), No co-expression of CD3, CD11b, CD13, CD14, CD16, CD20, CD33, CD64, CD117; B lymphoid blast population, Abnormal blast population 2: 11% of total events, Expressed variable CD15, CD33, variable CD34, CD38, weak CD71, CD123, Possible co-expression of weak-absent CD11b. | 46-47,XY,t(5;11)(q13;q23) | PTPN11 S502L (VAF 6.5%), NRAS G12D (VAF 12%), MPL F105L (VAF 48%) | Rituximab, cyclophosphamide, vincristine, doxorubicin, dexamethasone | CR | Y | 863 | Y |
| 5 | 76 | F | B/Myeloid | Expressed weak CD4, CD13, CD19, weak CD22, CD34, CD38, weak CD71, weak CD123, HLA-DR, Subset co-expressed CD117 (21% of gated events positive), without co-expression of CD22, Subset co-expressed CD10 (11% of gated events positive). | Complex: 45-46,XX,der(X;11)(q10;q10),add(2)(p11.2),del(5)(q31q33),i(8)(q10),del(11)(q13q23), der(15;22)(q10;q10),-16,add(20)(q11.2),+22,add(22)(p11.2)[cp19]/46,XX[1] | TP53 R248Q (VAF 96%), SF3B1 P580L (VAF 45%) | Inotuzumab, vincristine, prednisone, peg-asparaginase | Refractory disease | N | 51 | N |
| 6 | 70 | M | B/Myeloid | CD10+ CD13+ CD19+ cCD22+ CD25+ CD34+ cCD79a+ HLA-DR+ TdT+ (~38%); ** 2) CD4dim+ CD11b+ CD13+ CD14+ CD15+ CD33+ CD64+ HLA-DR(dim)+ (~30%). | 45,XY,-7,t(9;22)(q34;q11.2 | No mutations detected | Dasatinib, vincristine, dexamethasone, asparaginase, cytarabine | CR | N | 746 | N |
| 7 | 60 | F | B/Myeloid | Expressed weak CD13, weak CD15, weak CD33, CD34, CD38, variable CD64, weak CD71, variable CD117, weak CD123, HLA-DR, Variable expression of cytoplasmic CD22, CD7, Weak expression of TdT, Majority co-expressed CD19 (40% of total events positive), Subset co-expressed CD22 (20% of total events positive), Subset co-expressed MPO (6% of total events positive). | 46,XX,t(11;19)(q23;p13.1) | BRCA S1982fs (VAF 45%), DNMT3A A787T (VAF 44%), DNMT3A C537G (VAF 46%), IDH2 R140Q (VAF 7%) | Azacitidine, Venetoclax, Enasidenib | CRi | N | 333 | N |
| 8 | 56 | F | B/Myeloid | Expressed CD7, CD13, CD33, CD34, CD38, variable CD56, weak CD71, variable CD117, HLA-DR, Subset co-expressed CD15 (8% of total events positive), Subset co-expressed CD19 and CD22 (15% of total events positive) | Complex: 56-59<2n>,XX,+1,+8,+9,+10,+11,+11,+13,+14,+14,+19,+21,+22,+22 | TP53 S215G (VAF 80%), DNMT3A R882C (VAF 44%) | Daunorubicin, vincristine, prednisone, peg-asparaginase | Refractory Disease | Y | 371 | N |
| 9 | 54 | M | B/Myeloid | Expressed CD34, CD38, HLA-DR; Majority subset (88% of blasts): variable CD15, CD19, weak CD22, weak CD79a, Subset expressed TdT (8% of gated events positive); Minority subset (12% of blasts): weak CD11c, weak CD13, CD33, CD64 | Complex: 47,XY,+X,t(4;11)(q21;q23), del(17)(p11.2)[7]/48,idem,+8[3]/48~49,idem,+1,i(21)(q10),+mar[2]/46,XY[8]. | Not available | Daunorubicin, vincristine, prednisone, peg-asparaginase | Refractory Disease | N | 132 | N |

HCT: hematopoietic cell transplantation; CR: Complete response; CRi: Complete response with incomplete hematologic recovery; VAF: variant allele frequency

| Patient Number | Age at MPAL Diagnosis | Sex | Clinical Subtype | Clinical Flow Cytometry | Cytogenetics | Clinical Bulk Next Generation Sequencing | Frontline Therapy | Clinical response to frontline therapy | Allogeneic HCT? | Survival After MPAL Diagnosis (days) | Alive? |
|---|---|---|---|---|---|---|---|---|---|---|---|
| 10 | 53 | F | B/Myeloid | Expressed CD10(small subset)+, CD11b(small subset)+, CD13+, CD14(very small subset)+, CD15 (small subset)+, CD19(large subset)+, CD33+, CD34(subset)+, CD64(subset)+, HLA-DR+, MPO(dim)+, TdT(subset)+, cCD79a(subset)+. | Not available | Not available | Daunorubicin and cytarabine | CR | Y | 5403 | N |
| 11 | 59 | F | T/Myeloid | Expressed CD2, CD4, CD7, CD13, weak-absent CD33, CD34, CD38, weak CD71, CD117, HLA-DR, Some possible co-expression of weak-absent CD123, Cytoplasmic expression of variable CD3, No co-expression of CD14, CD15, CD16, MPO, TdT, B-cell antigens | Normal female karyotype | Not available | Daunorubicin, vincristine, prednisone, peg-asparaginase | CRi | N | 353 | N |
| 12 | 80 | F | T/Myeloid | Expressed: cCD3(predominantly)+, CD5 (dim)+ CD7+ CD10(var)+ CD33 (var)+ CD34(predominantly)+ CD38+ HLA-DR(var)+ MPO(subset)+ TdT- CD79a-. | Normal female karyotype | NRAS G12C (VAF 48%), DNMT3A W306G (VAF 49%), IDH1 R132C (VAF 48%), ETV6 F287fs (VAF 45%), NOTCH1 V1605ins (VAF 32%), NOTCH1 A1701P (VAF 11%) | Vincristine and prednisone | N/A | N | 64 | N |
| 13 | 80 | F | T/Myeloid | Expression demonstrates two discrete populations of blasts, each accounting for 40% of total cellularity: (1) A population with low CD45/side scatter and cCD3(subset)+ CD5+ CD7(bright)+ CD13- CD33(bright)+ CD34+ HLA-DR- MPO- TdT(subset)+. (2) A population with intermediate CD45/side scatter and CD2(dim)+ CD11b+ CD13+ CD14+ CD15- CD33+ CD34- CD64+ HLA-DR(variable)+ MPO+ TdT-. | 45,XX,-7,del(17)t(7;17)(p11.2;p11.2) | Not available | Cytarabine | Refractory Disease | N | 84 | N |
| 14 | 43 | F | B and T/Myeloid | Expressed weak CD2, bright CD7, weak CD13, variable CD33, CD34, CD38, weak CD71, weak-absent CD117, weak-absent HLA-DR; B lymphoid blasts (4%): CD10, CD19, weak CD20, weak CD22, weak CD33, CD34, CD38, HLA-DR | Normal female karyotype | DNMT3A T536* (VAF 45.6%), DNMT3A C562T (VAF 43%), IDH2 R140Q (VAF 45%), PHF6 R116* (VAF 45%), NRAS G12D (VAF 46%), NOTCH1 F1592S (VAF 49%), EZH2 T731H (VAF 42%), EZH2 H507R (VAF 2.2%) | Daunorubicin, vincristine, prednisone, peg-asparaginase | Refractory Disease | Y | 182 | N |

HCT: hematopoietic cell transplantation; CR: Complete response; CRi: Complete response with incomplete hematologic recovery; VAF: variant allele frequency

### Supplementary Table 2: Single-Cell DAb-seq Amplicon Panel

| Chromosome | Gene | Amplicon Start | Amplicon End |
|---|---|---|---|
| chr1 | NRAS | 115256488 | 115256723 |
| chr1 | NRAS | 115258610 | 115258825 |
| chr2 | DNMT3A | 25457144 | 25457372 |
| chr2 | DNMT3A | 25458519 | 25458763 |
| chr2 | DNMT3A | 25459794 | 25460046 |
| chr2 | DNMT3A | 25461881 | 25462137 |
| chr2 | DNMT3A | 25463107 | 25463346 |
| chr2 | DNMT3A | 25463494 | 25463717 |
| chr2 | DNMT3A | 25464422 | 25464618 |
| chr2 | DNMT3A | 25466622 | 25466871 |
| chr2 | DNMT3A | 25467013 | 25467220 |
| chr2 | DNMT3A | 25467372 | 25467631 |
| chr2 | DNMT3A | 25468111 | 25468332 |
| chr2 | DNMT3A | 25469007 | 25469231 |
| chr2 | DNMT3A | 25469408 | 25469606 |
| chr2 | DNMT3A | 25469926 | 25470185 |
| chr2 | DNMT3A | 25470404 | 25470663 |
| chr2 | DNMT3A | 25470931 | 25471190 |
| chr2 | DNMT3A | 25472516 | 25472728 |
| chr2 | SF3B1 | 198266104 | 198266322 |
| chr2 | SF3B1 | 198266443 | 198266688 |
| chr2 | SF3B1 | 198266694 | 198266913 |
| chr2 | SF3B1 | 198267320 | 198267569 |
| chr2 | IDH1 | 209113086 | 209113297 |
| chr3 | GATA2 | 128200078 | 128200327 |
| chr3 | GATA2 | 128200669 | 128200928 |
| chr3 | GATA2 | 128202700 | 128202899 |
| chr4 | KIT | 55561580 | 55561792 |
| chr4 | KIT | 55569876 | 55570095 |
| chr4 | KIT | 55592078 | 55592282 |
| chr4 | KIT | 55593510 | 55593744 |
| chr4 | KIT | 55593964 | 55594183 |
| chr4 | KIT | 55599271 | 55599486 |
| chr4 | KIT | 55602652 | 55602862 |
| chr4 | TET2 | 106154924 | 106155158 |
| chr4 | TET2 | 106155159 | 106155416 |
| chr4 | TET2 | 106155470 | 106155729 |
| chr4 | TET2 | 106155914 | 106156173 |
| chr4 | TET2 | 106156238 | 106156489 |
| chr4 | TET2 | 106156504 | 106156762 |
| chr4 | TET2 | 106156794 | 106157046 |
| chr4 | TET2 | 106157078 | 106157332 |
| chr4 | TET2 | 106157427 | 106157679 |
| chr4 | TET2 | 106157757 | 106158015 |
| chr4 | TET2 | 106158030 | 106158288 |
| chr4 | TET2 | 106158294 | 106158544 |
| chr4 | TET2 | 106158546 | 106158805 |
| chr4 | TET2 | 106162389 | 106162618 |
| chr4 | TET2 | 106163953 | 106164192 |
| chr4 | TET2 | 106164706 | 106164952 |
| chr4 | TET2 | 106180719 | 106180956 |
| chr4 | TET2 | 106182830 | 106183072 |
| chr4 | TET2 | 106190734 | 106190955 |
| chr4 | TET2 | 106193541 | 106193777 |
| chr4 | TET2 | 106193778 | 106194032 |
| chr4 | TET2 | 106194036 | 106194295 |
| chr4 | TET2 | 106196180 | 106196429 |
| chr4 | TET2 | 106196438 | 106196692 |
| chr4 | TET2 | 106196772 | 106197024 |
| chr4 | TET2 | 106197029 | 106197278 |
| chr4 | TET2 | 106197336 | 106197593 |
| chr4 | TET2 | 106197599 | 106197858 |
| chr5 | NPM1 | 170837385 | 170837659 |

| Chromosome | Gene | Amplicon Start | Amplicon End |
|---|---|---|---|
| chr7 | EZH2 | 148504722 | 148504971 |
| chr7 | EZH2 | 148506026 | 148506265 |
| chr7 | EZH2 | 148506372 | 148506589 |
| chr7 | EZH2 | 148507405 | 148507618 |
| chr7 | EZH2 | 148508697 | 148508930 |
| chr7 | EZH2 | 148511033 | 148511276 |
| chr7 | EZH2 | 148511991 | 148512230 |
| chr7 | EZH2 | 148514918 | 148515124 |
| chr7 | EZH2 | 148523627 | 148523866 |
| chr7 | EZH2 | 148525654 | 148525888 |
| chr7 | EZH2 | 148526738 | 148526948 |
| chr7 | EZH2 | 148529631 | 148529890 |
| chr7 | EZH2 | 148543468 | 148543693 |
| chr7 | EZH2 | 148544267 | 148544493 |
| chr9 | JAK2 | 5073699 | 5073902 |
| chr11 | WT1 | 32413428 | 32413633 |
| chr11 | WT1 | 32414186 | 32414405 |
| chr11 | WT1 | 32417759 | 32417989 |
| chr11 | WT1 | 32421512 | 32421750 |
| chr11 | WT1 | 32439083 | 32439321 |
| chr12 | KRAS | 25378535 | 25378794 |
| chr12 | KRAS | 25380239 | 25380478 |
| chr12 | KRAS | 25398207 | 25398433 |
| chr12 | PTPN11 | 112888116 | 112888350 |
| chr12 | PTPN11 | 112890995 | 112891234 |
| chr12 | PTPN11 | 112910668 | 112910907 |
| chr12 | PTPN11 | 112915378 | 112915582 |
| chr12 | PTPN11 | 112924189 | 112924405 |
| chr12 | PTPN11 | 112926204 | 112926424 |
| chr12 | PTPN11 | 112926825 | 112927050 |
| chr13 | FLT3 | 28589757 | 28589977 |
| chr13 | FLT3 | 28592474 | 28592726 |
| chr13 | FLT3 | 28597498 | 28597727 |
| chr13 | FLT3 | 28601131 | 28601358 |
| chr13 | FLT3 | 28602303 | 28602559 |
| chr13 | FLT3 | 28608189 | 28608395 |
| chr13 | FLT3 | 28608474 | 28608696 |
| chr13 | FLT3 | 28609572 | 28609790 |
| chr13 | FLT3 | 28610015 | 28610260 |
| chr15 | IDH2 | 90631741 | 90631990 |
| chr17 | TP53 | 7572907 | 7573129 |
| chr17 | TP53 | 7573974 | 7574178 |
| chr17 | TP53 | 7576760 | 7576976 |
| chr17 | TP53 | 7577015 | 7577264 |
| chr17 | TP53 | 7577398 | 7577636 |
| chr17 | TP53 | 7578076 | 7578315 |
| chr17 | TP53 | 7578363 | 7578618 |
| chr17 | TP53 | 7579859 | 7580118 |
| chr17 | SRSF2 | 74732192 | 74732450 |
| chr20 | ASXL1 | 30956750 | 30956969 |
| chr20 | ASXL1 | 31015814 | 31016051 |
| chr20 | ASXL1 | 31021138 | 31021366 |
| chr20 | ASXL1 | 31021439 | 31021659 |
| chr20 | ASXL1 | 31022169 | 31022417 |
| chr20 | ASXL1 | 31022720 | 31022978 |
| chr20 | ASXL1 | 31023009 | 31023248 |
| chr20 | ASXL1 | 31023262 | 31023492 |
| chr20 | ASXL1 | 31023557 | 31023761 |
| chr21 | RUNX1 | 36171568 | 36171811 |
| chr21 | RUNX1 | 36206684 | 36206913 |
| chr21 | RUNX1 | 36231693 | 36231937 |
| chr21 | RUNX1 | 36252820 | 36253046 |
| chr21 | U2AF1 | 44514550 | 44514808 |
| chr21 | U2AF1 | 44524417 | 44524634 |

### Supplementary Table 3: Antibody-Oligo Conjugates (AOCs) for Single Cell DAb-seq

| Product Name | barcode | Clone | Supplier Link |
|---|---|---|---|
| CD3 | AACGCTTC | UCHT1 | https://www.biolegend.com/en-us/products/ultra-leaf-purified-anti-human-cd3-antibody-7742 |
| CD4 | CGGTTACA | OKT4 | https://www.biolegend.com/en-us/products/leaf-low-endotoxin--azide-freepurified-anti-human-cd4-antibody-3651 |
| CD5 | CCACTTAG | UCHT2 | https://www.miltenyibiotec.com/US-en/products/macs-flow-cytometry/antibodies/primary-antibodies/cd5-antibodies-human-ucht2.html#pure:100-ug-in-100-ul |
| CD7 | GCCAAGTT | 848438 | https://www.rndsystems.com/products/human-cd7-antibody-848438_mab7579 |
| CD8 | CGACAAGA | REA734 | https://www.miltenyibiotec.com/US-en/products/macs-flow-cytometry/antibodies/primary-antibodies/cd8-antibodies-human-rea734.html#pure:100-ug-in-100-ul |
| CD10 | TGGCAGAA | 97C5 | https://www.miltenyibiotec.com/US-en/products/macs-flow-cytometry/antibodies/primary-antibodies/cd10-antibodies-human-97c5.html#pure:100-ug-in-100-ul |
| CD11b | ATGTAGCC | ICRF44 | https://www.biolegend.com/en-us/products/leaf-low-endotoxin--azide-freepurified-anti-human-cd11b-antibody-767 |
| CD13 | ACGGAATG | WM15 | https://www.biolegend.com/en-us/products/ultra-leaf-purified-anti-human-cd13-antibody-15458 |
| CD14 | TGTGACGT | M5E2 | https://www.biolegend.com/en-us/products/leaf-low-endotoxin--azide-freepurified-anti-human-cd14-antibody-795 |
| CD15 | AACCGAGA | ICRF29-2 | https://www.rndsystems.com/products/human-cd15-lewis-x-antibody-icrf29-2_mab7368 |
| CD16 | ACAAGGAC | 3G8 | https://www.biolegend.com/en-us/products/ultra-leaf-purified-anti-human-cd16-antibody-8089 |
| CD19 | ACTGCCAA | HIB19 | https://www.biolegend.com/en-us/products/leaf-low-endotoxin--azide-freepurified-anti-human-cd19-antibody-718 |
| CD22 | AAGGTGGT | 219934 | https://www.rndsystems.com/products/human-siglec-2-cd22-antibody-219934_mab1968 |
| CD30 | AGGTCCTA | 81337 | https://www.rndsystems.com/products/human-cd30-tnfrsf8-antibody-81337_mab229 |
| CD33 | GGAACCAT | AC104.3E3 | https://www.miltenyibiotec.com/US-en/products/macs-flow-cytometry/antibodies/primary-antibodies/cd33-antibodies-human-ac104-3e3.html#pure:100-ug-in-100-ul |
| CD34 | CAGAGCTA | AC136 | https://www.miltenyibiotec.com/US-en/products/macs-flow-cytometry/antibodies/primary-antibodies/cd34-antibodies-human-ac136.html#pure:100-ug-in-100-ul |
| CD38 | ACCTCACT | REA671 | https://www.miltenyibiotec.com/US-en/products/macs-flow-cytometry/antibodies/primary-antibodies/cd38-antibodies-human-rea671.html#pure:100-ug-in-100-ul |
| CD56 | CCTTGATC | REA196 | https://www.miltenyibiotec.com/US-en/products/macs-flow-cytometry/antibodies/primary-antibodies/cd56-antibodies-human-rea196.html#pure:100-ug-in-100-ul |
| CD64 | AACAACCG | 10.1.1 | https://www.miltenyibiotec.com/US-en/products/macs-flow-cytometry/antibodies/primary-antibodies/cd64-antibodies-human-10-1-1.html#pure:100-ug-in-100-ul |
| CD71 | AGATTCGC | AC102 | https://www.miltenyibiotec.com/US-en/products/macs-flow-cytometry/antibodies/primary-antibodies/cd71-antibodies-human-ac102.html#pure:100-ug-in-100-ul |
| CD117 | TTCGTTGG | HLDA6 | https://www.biolegend.com/en-us/products/leaf-low-endotoxin--azide-freepurified-anti-human-cd117-c-kit-antibody-3716 |
| CD123 | GATGGTCA | AC145 | https://www.miltenyibiotec.com/US-en/products/macs-flow-cytometry/antibodies/primary-antibodies/cd123-antibodies-human-ac145.html#pure:100-ug-in-100-ul |
| CD45-26 | AGTGGTAC | 5B1 | https://www.miltenyibiotec.com/US-en/products/macs-flow-cytometry/antibodies/primary-antibodies/cd45-antibodies-human-5b1.html#pure:100-ug-in-100-ul |
| CD45-27 | ATTGGCTG | 5B1 | https://www.miltenyibiotec.com/US-en/products/macs-flow-cytometry/antibodies/primary-antibodies/cd45-antibodies-human-5b1.html#pure:100-ug-in-100-ul |
| CD45-28 | CAGTCGAA | 5B1 | https://www.miltenyibiotec.com/US-en/products/macs-flow-cytometry/antibodies/primary-antibodies/cd45-antibodies-human-5b1.html#pure:100-ug-in-100-ul |

Supplementary Table 4: Number of Single Cells Isolated per Patient per Assay

| Patient Number | Single Cell Assay Used | DNA + Protein: Number of Single Cells Isolated | RNA + Protein: Number of Single Cells Isolated |
|---|---|---|---|
| 1 | DNA+protein, RNA+ protein | 4274 | 9386 |
| 2 | DNA+protein, RNA+ protein | 1093 | 6100 |
| 3 | DNA+protein, RNA+ protein | 3911 | 9649 |
| 4 | DNA+protein, RNA+ protein | 2848 | 7395 |
| 5 | DNA+protein, RNA+ protein | 4649 | 5494 |
| 6 | DNA+protein, RNA+ protein | 5571 | 1173 |
| 7 | DNA+protein, RNA+ protein | 3339 | 2578 |
| 8 | DNA+protein, RNA+ protein | 4895 | 9012 |
| 9 | DNA+protein, RNA+ protein | 2632 | 6252 |
| 10 | DNA+protein only | 1660 | - |
| 11 | DNA+protein, RNA+ protein | 5472 | 2594 |
| 12 | DNA+protein, RNA+ protein | 7245 | 10275 |
| 13 | DNA+protein, RNA+ protein | 7136 | - |
| 14 | DNA+protein, RNA+ protein | 4082 | 2223 |

### Supplementary Table 5: Variants and Variant Allele Frequencies Called by SC DAb-seq

| Patient Number | Variant | Protein | SC VAF (Cell Count) | SC VAF (Read Count) |
|---|---|---|---|---|
| 1 | TP53:chr17:7578190:T/C | Y220C | 23.3% Het; 23.7% Hom | 24.5% Het; 24.8% Hom. |
| 1 | JAK2:chr9:5073770:G/T | V617F | 0.51% | 0.46% |
| 2 | DNMT3A:chr2:25457243:G/A | R882C | 2.30% | 3.90% |
| 2 | KRAS:chr12:25398284:C/T | G12D | 7.90% | 11.40% |
| 3 | FLT3:chr13:28608301:T/TCAACGTA' | ITD (c.1781) | 41.90% | 42.00% |
| 3 | FLT3:chr13:28608302:G/GCCAAATCAACGT | ITD (c.1813) | 6.50% | 7.60% |
| 3 | SF3B1:chr2:198267361:T/C | K666E | 78% Het; 3% Hom | 72% Het; 5.2% Hom |
| 3 | PTPN11:chr12:112888199:C/T | A72V | 0.62% | 0.49% |
| 4 | NRAS:chr1:115258747:C/T | G12D | 14.40% | 13.60% |
| 4 | PTPN11:chr12:112926885:C/T | S502L | 7.80% | 7.00% |
| 4 | RUNX1:chr21:36206776:T/G | T246fs | 2.40% | 2.10% |
| 4 | RUNX1:chr21:36206770:T/G | N248H | 1.60% | 1.40% |
| 4 | TP53:chr17:7578385:C/T' | C182Y | 0.52% | 0.46% |
| 4 | U2AF1:chr21:44524472:G/T | H29N | 0.61% | 0.48% |
| 5 | TP53:chr17:7577538:C/T' | R248Q | 9.5% Het; 83.8% Hom | 12.1% Het; 85.9% Hom |
| 6 | RUNX1:chr21:36206776:T/G | T246P | 2.50% | 2.10% |
| 6 | EZH2:chr7:148504716:AG/A | EZH2 c.*21del | 11.90% | 9.10% |
| 7 | IDH2:chr15:90631934:C/T | R140Q | 24% | 26.40% |
| 7 | FLT3:chr13:28592642:C/A | D835Y | 0.80% | 2.20% |
| 7 | GATA2:chr3:128202770:T/C | N317S | 0.50% | 1.20% |
| 8 | DNMT3A:chr2:25457243:G/A | R288C | 86.50% | 79.90% |
| 8 | TP53:chr17:7578206:T/C | S215G | 3.6% Het.; 80.0% Hom. | 2.4% Het.; 78.8% Hom. |
| 9 | TP53:chr17:7578210:T/C | R213= | 0.264 | 0.228 |
| 10 | Not mutations detected. | | | |
| 11 | FLT3:chr13:28602226:A/AGAGAGAGAGAGAGAG | ITD (c.1802) | 83.50% | 86.10% |
| 11 | DNMT3A:chr2:25457243:G/A | R288C | 82.40% | 87.90% |
| 11 | ASXL1:chr20:31022910:G/T | D799Y | 84.10% | 94.10% |
| 11 | RUNX1:chr21:36252869:C/CGGGGCCCCAT | G165fs | 0.92% | 1.10% |
| 12 | NRAS:chr1:115258748:C/T' | G12S | 72.9% Het; 2.8% Hom. | 78.6% Het; 3.6% Hom. |
| 12 | DNMT3A:chr2:25457242:C/T | W306G | 14.30% | 15.60% |
| 12 | IDH1:chr2:209113113:G/A | R132C | 74.00% | 74.30% |
| 13 | DNMT3A:chr2:25457242:C/T | R288C | 95.70% | 92.00% |
| 14 | NRAS:chr1:115258747:C/T | G12D | 85% Het.; 2.3% Hom. | 82.4% Het.; 1.9% Hom. |
| 14 | NRAS:chr1:115258748:C/T' | G12S | 3.70% | 2.80% |
| 14 | IDH2:chr15:90631934:C/T | R140Q | 83.50% | 79.80% |
| 14 | IDH2:chr15:90631917:TC/T | T146fs | 0.60% | 0.50% |
| 14 | U2AF1:chr21:44524472:G/T | H29N | 25% | 22.10% |

Supplementary Table 6: Antibody-Oligo Conjugates (AOCs) for Single Cell CITE-seq

| Name | Barcode | Description | BioLegend Cat. No. | Clone |
|---|---|---|---|---|
| ADT-CD14 | TCTCAGACCTCCGTA | TotalSeq™-A0081 anti-human CD14 Antibody | 301855 | M5E2 |
| ADT-CD64 | AAGTATGCCCTACGA | TotalSeq™-A0162 anti-human CD64 Antibody | 305037 | 10.1 |
| ADT-CD22 | GGGTTGTTGTCTTTG | TotalSeq™-A0393 anti-human CD22 Antibody | 363514 | S-HCL-1 |
| ADT-CD10 | CAGCCATTCATTAGG | TotalSeq™-A0062 anti-human CD10 Antibody | 312231 | HI10a |
| ADT-CD13 | TTTCAACGCCCTTTC | TotalSeq™-A0364 anti-human CD13 Antibody | 301729 | WM15 |
| ADT-CD117 | AGACTAATAGCTGAC | TotalSeq™-A0061 anti-human CD117 (c-kit) Antibody | 313241 | 104D2 |
| ADT-CD5 | CATTAACGGGATGCC | TotalSeq™-A0138 anti-human CD5 Antibody | 300635 | UCHT2 |
| ADT-CD7 | TGGATTCCCGGACTT | TotalSeq™-A0066 anti-human CD7 Antibody | 343123 | CD7-6B7 |
| ADT-CD56 | TTCGCCGCATTGAGT | TotalSeq™-A0084 anti-human CD56 (NCAM) Recombinant Antibody | 392421 | QA17A16 |
| ADT-HLA-DR | AATAGCGAGCAAGTA | TotalSeq™-A0159 anti-human HLA-DR Antibody | 307659 | L243 |
| ADT-CD11b | GACAAGTGATCTGCA | TotalSeq™-A0161 anti-human CD11b Antibody | 301353 | ICRF44 |
| ADT-CD4 | GCGATCCCTTGAGAT | TotalSeq™-A0922 anti-human CD4 Antibody | 317451 | OKT4 |
| ADT-IgG1 | GCCGGACGACATTAA | TotalSeq™-A0090 Mouse IgG1, κ isotype Ctrl Antibody | 400199 | MOPC-21 |
| ADT-CD3 | TATCCCTTGGGATGG | TotalSeq™-A0049 anti-human CD3 Antibody | 344847 | SK7 |
| ADT-CD19 | CTGGGCAATTACTCG | TotalSeq™-A0050 anti-human CD19 Antibody | 302259 | HIB19 |
| ADT-CD30 | TCAGGGTGTGCTGTA | TotalSeq™-A0028 anti-human CD30 Antibody | 333913 | BY88 |
| ADT-CD33 | TAACTCAGGGCCTAT | TotalSeq™-A0052 anti-human CD33 Antibody | 366629 | P67.6 |
| ADT-CD45 | TGCAATTACCCGGAT | TotalSeq™-A0391 anti-human CD45 Antibody | 304064 | HI30 |
| ADT-CD34 | GCAGAAATCTCCCTT | TotalSeq™-A0054 anti-human CD34 Antibody | 343537 | 581 |

### Supplementary Table 7: Transcriptional Signature of Conserved Leukemia Signature Across 12 Patients with MPAL

| | Patient 1 | | Patient 2 | | Patient 3 | | Patient 4 | |
|---|---|---|---|---|---|---|---|---|
| | Average Log2 Fold Change | Adjusted p-value | Average Log2 Fold Change | Adjusted p-value | Average Log2 Fold Change | Adjusted p-value | Average Log2 Fold Change | Adjusted p-value |
| RPLP0 | 0.373275671 | 3.01E-42 | 1.821453275 | 0 | 0.883831482 | 0 | 0.749040469 | 1.07E-93 |
| DNTT | 1.966830284 | 0 | 1.888098601 | 4.33E-125 | 1.913390095 | 0 | 1.803757004 | 1.03E-186 |
| H2AFY | 1.383998651 | 4.56E-294 | 0.96544454 | 6.69E-56 | 0.537731359 | 6.47E-30 | 1.494398295 | 1.73E-201 |
| HNRNPA1 | 0.611397698 | 2.24E-118 | 1.307375476 | 1.51E-217 | 0.563950016 | 1.26E-71 | 0.539508058 | 2.11E-69 |
| HMGA1 | 1.150623978 | 4.56E-155 | 2.220053497 | 1.79E-220 | 0.719815173 | 1.64E-44 | 1.545766103 | 2.53E-167 |
| TUBB | 1.345920531 | 4.16E-181 | 2.066641109 | 3.55E-132 | 0.924220579 | 3.21E-91 | 1.855080792 | 2.54E-173 |
| ENO1 | 1.045671096 | 2.47E-169 | 0.742864314 | 2.89E-47 | 0.599110681 | 1.20E-57 | 0.558525523 | 3.61E-42 |
| MSI2 | 1.172336703 | 2.99E-169 | 0.459647993 | 1.97E-15 | 1.054383605 | 2.75E-107 | 0.970979939 | 3.58E-74 |
| IMPDH2 | 0.579197577 | 2.68E-57 | 1.794577424 | 3.17E-144 | 1.057081929 | 4.47E-100 | 1.148924194 | 8.63E-115 |
| TRIM28 | 0.802245453 | 3.37E-116 | 0.979203772 | 3.16E-61 | 0.812223076 | 1.30E-55 | 0.844841442 | 1.94E-79 |
| PARP1 | 0.73949727 | 1.52E-99 | 0.982515031 | 1.96E-57 | 0.36301667 | 1.79E-14 | 0.727247059 | 1.40E-65 |
| CHD4 | 0.636451283 | 3.18E-69 | 0.717214962 | 8.31E-31 | 0.654458878 | 1.09E-38 | 0.976532872 | 6.76E-91 |

| | Patient 5 | | Patient 6 | | Patient 7 | | Patient 8 | |
|---|---|---|---|---|---|---|---|---|
| | Average Log2 Fold Change | Adjusted p-value | Average Log2 Fold Change | Adjusted p-value | Average Log2 Fold Change | Adjusted p-value | Average Log2 Fold Change | Adjusted p-value |
| RPLP0 | 1.118454185 | 2.40E-75 | 0.377010072 | 1 | 0.91662864 | 5.21E-45 | 0.342569995 | 3.05E-83 |
| DNTT | 1.661978611 | 2.99E-25 | 0.66247969 | 1 | 1.656084368 | 9.07E-37 | 0.950392327 | 8.44E-43 |
| H2AFY | 0.828313697 | 1.00E-05 | 0.456674459 | 1 | 1.678310079 | 4.97E-69 | 0.576939745 | 7.05E-65 |
| HNRNPA1 | 0.526683424 | 2.95E-06 | 0.466270117 | 1 | 0.992495303 | 9.20E-52 | 0.512461328 | 3.21E-87 |
| HMGA1 | 1.224370787 | 9.62E-10 | 0.494420964 | 1 | 1.117224107 | 2.15E-21 | 0.436011044 | 4.60E-13 |
| TUBB | 0.886887209 | 2.81E-09 | 0.418751211 | 1 | 1.939951043 | 1.93E-56 | 0.818113862 | 2.73E-63 |
| ENO1 | 0.986972506 | 1.69E-17 | 0.548580586 | 1 | 1.117043022 | 1.86E-40 | 0.816215094 | 8.12E-147 |
| MSI2 | 0.435445685 | 1 | 0.483872436 | 1 | 1.155202672 | 2.60E-22 | 0.415159503 | 2.46E-21 |
| IMPDH2 | 0.529149495 | 0.291423744 | 0.862650739 | 1 | 0.928880815 | 1.56E-21 | 0.89812216 | 1.34E-78 |
| TRIM28 | 0.544268506 | 0.105908817 | 0.491345784 | 1 | 0.739034007 | 2.02E-15 | 0.520527217 | 1.10E-29 |
| PARP1 | 0.818476176 | 4.13E-07 | 0.307368011 | 1 | 0.699686417 | 1.87E-14 | 0.592702544 | 3.51E-33 |
| CHD4 | 0.314126946 | 1 | 0.499785279 | 1 | 0.590530033 | 5.63E-07 | 0.376670844 | 3.42E-13 |

| | Patient 9 | | Patient 11 | | Patient 12 | | Patient 14 | |
|---|---|---|---|---|---|---|---|---|
| | Average Log2 Fold Change | Adjusted p-value | Average Log2 Fold Change | Adjusted p-value | Average Log2 Fold Change | Adjusted p-value | Average Log2 Fold Change | Adjusted p-value |
| RPLP0 | 0.970656145 | 3.36E-122 | 0.763630696 | 1.45E-60 | 0.628921105 | 5.89E-240 | 1.0332466 | 1.65E-32 |
| DNTT | 1.389285995 | 2.66E-30 | 0.972662076 | 2.17E-07 | 1.549061452 | 1.48E-210 | 1.936191925 | 1.18E-14 |
| H2AFY | 0.787574792 | 4.11E-14 | 0.259561635 | 0.287406512 | 1.052907325 | 1.85E-216 | 1.367785701 | 2.69E-26 |
| HNRNPA1 | 0.266819422 | 0.066954507 | 0.831048838 | 3.69E-39 | 0.788802984 | 5.89E-231 | 1.361381513 | 3.27E-41 |
| HMGA1 | 1.377531919 | 2.08E-50 | 0.338362343 | 1 | 0.348917238 | 7.71E-28 | 0.759469519 | 0.001475125 |
| TUBB | 1.002707057 | 4.52E-27 | 1.329161753 | 1.92E-19 | 0.981960838 | 5.41E-142 | 1.063651921 | 1.79E-05 |
| ENO1 | 0.43863873 | 1.74E-10 | 0.925781334 | 1.87E-23 | 0.88468916 | 3.15E-159 | 1.363871326 | 8.64E-25 |
| MSI2 | 0.716557122 | 4.53E-10 | 0.598648985 | 0.001638696 | 0.649431083 | 1.17E-87 | 0.831780282 | 0.134162329 |
| IMPDH2 | 0.959426915 | 2.83E-20 | 1.004571549 | 2.35E-16 | 0.892602039 | 3.38E-155 | 1.028340521 | 1.13E-07 |
| TRIM28 | 0.927563585 | 2.94E-32 | 0.40879403 | 0.075438318 | 0.315887431 | 5.13E-35 | 1.0220468 | 2.98E-08 |
| PARP1 | 0.603335486 | 1.00E-06 | 0.424105636 | 0.072005089 | 0.263914272 | 8.98E-22 | 0.383760863 | 1 |
| CHD4 | 0.693025444 | 1.66E-10 | 0.370975804 | 1 | 0.374660522 | 1.26E-37 | 0.716564796 | 0.033794212 |

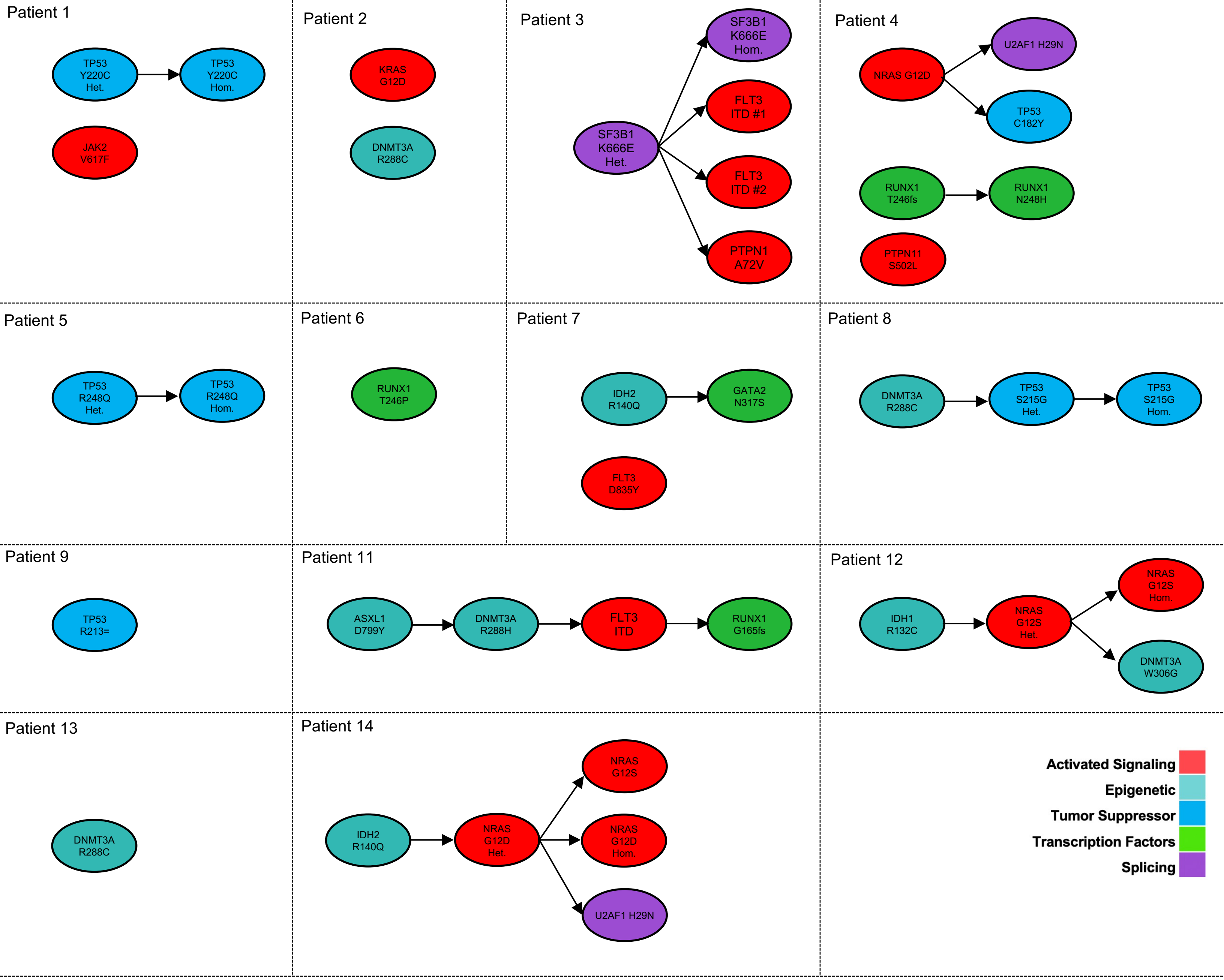


**Supplementary Figure 1.** Mutation phylogeny of 14 patients with newly-diagnosed MPAL derived from single-cell DNA sequencing using the SCITE algorithm. Each oval represents a subclone and arrows represent cumulative acquisition of mutational events. Subclones are color-coded based on biologic function. Het: Heterozygous; Hom: Homozygous. All mutations are heterozygous unless specified otherwise. Note Patient 10 had no detectable mutations.

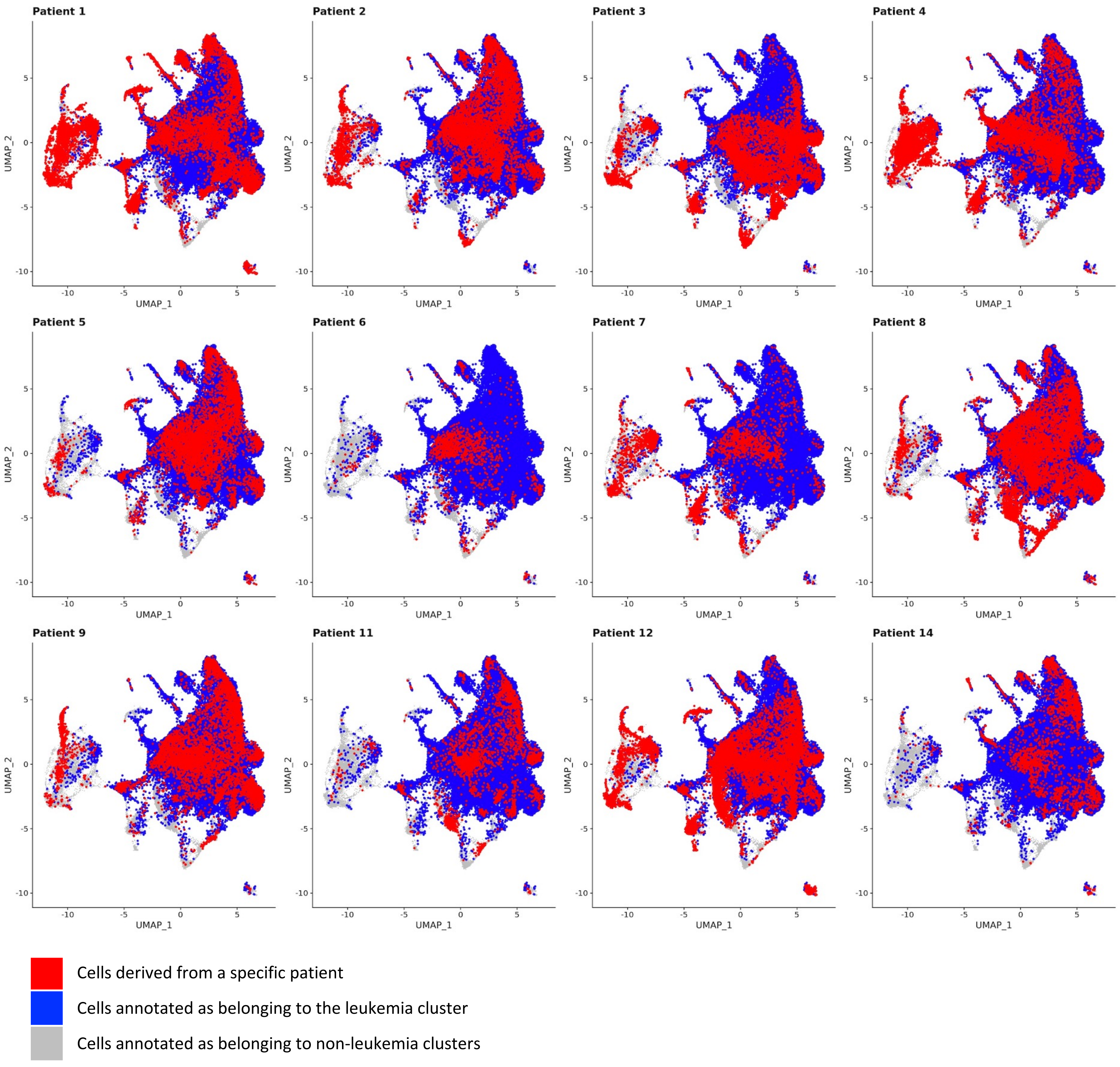


**Supplementary Figure 2**. RNA-derived UMAP from SC RNA+protein analysis of 71,579 cells from 12 patients. For each of the 12 panels, cells color-coded in red are derived from each of the 12 patients, cells color-coded in blue are in the leukemia cluster, and cells color-coded in grey are in non-leukemia clusters. All patients contributed to the leukemia cluster.

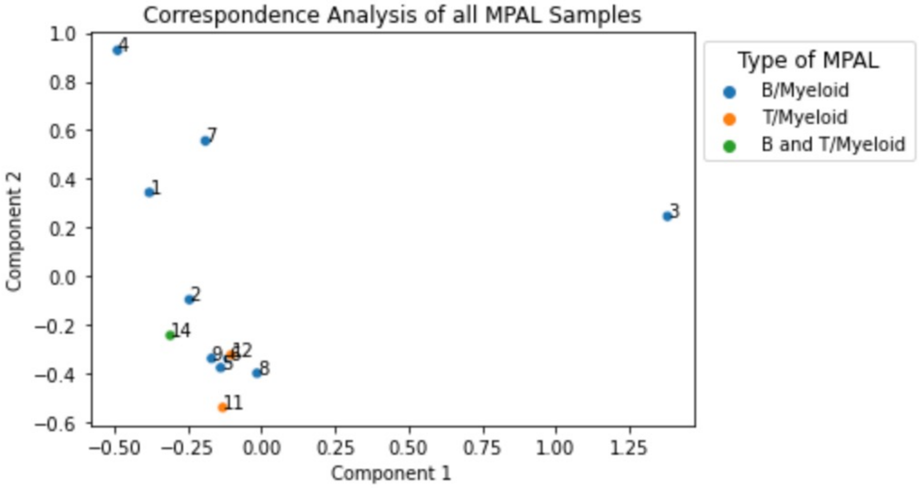


**Supplementary Figure 3.** Correspondence Analysis based on transcriptional data from 12 patients with MPAL. Each point represents an individual patient with patient numbers overlain. Points are color-coded based on immunophenotypic subtype.

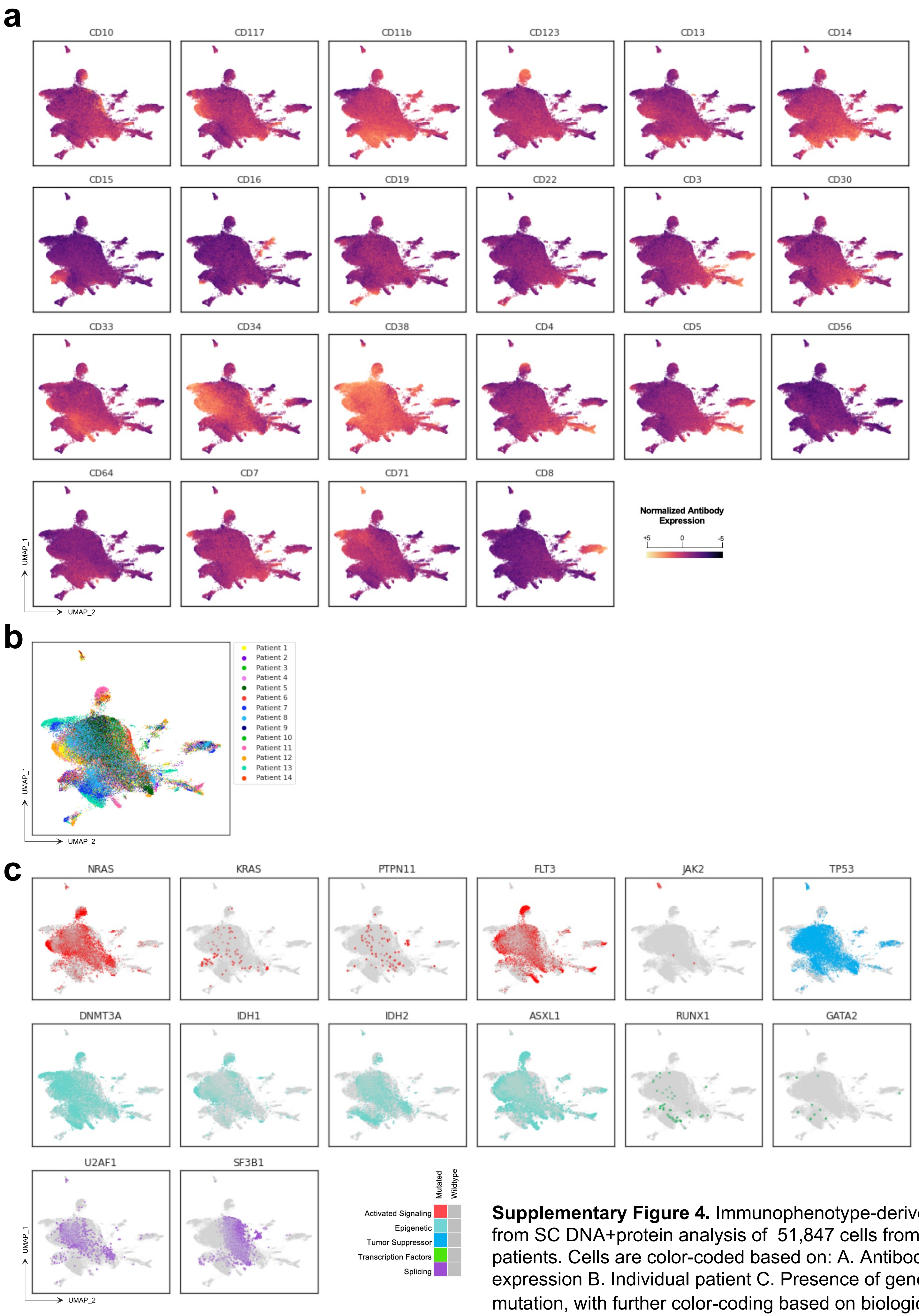


**Supplementary Figure 4.** Immunophenotype-derived UMAP from SC DNA+protein analysis of 51,847 cells from 14 patients. Cells are color-coded based on: A. Antibody expression B. Individual patient C. Presence of genetic mutation, with further color-coding based on biological function.

a

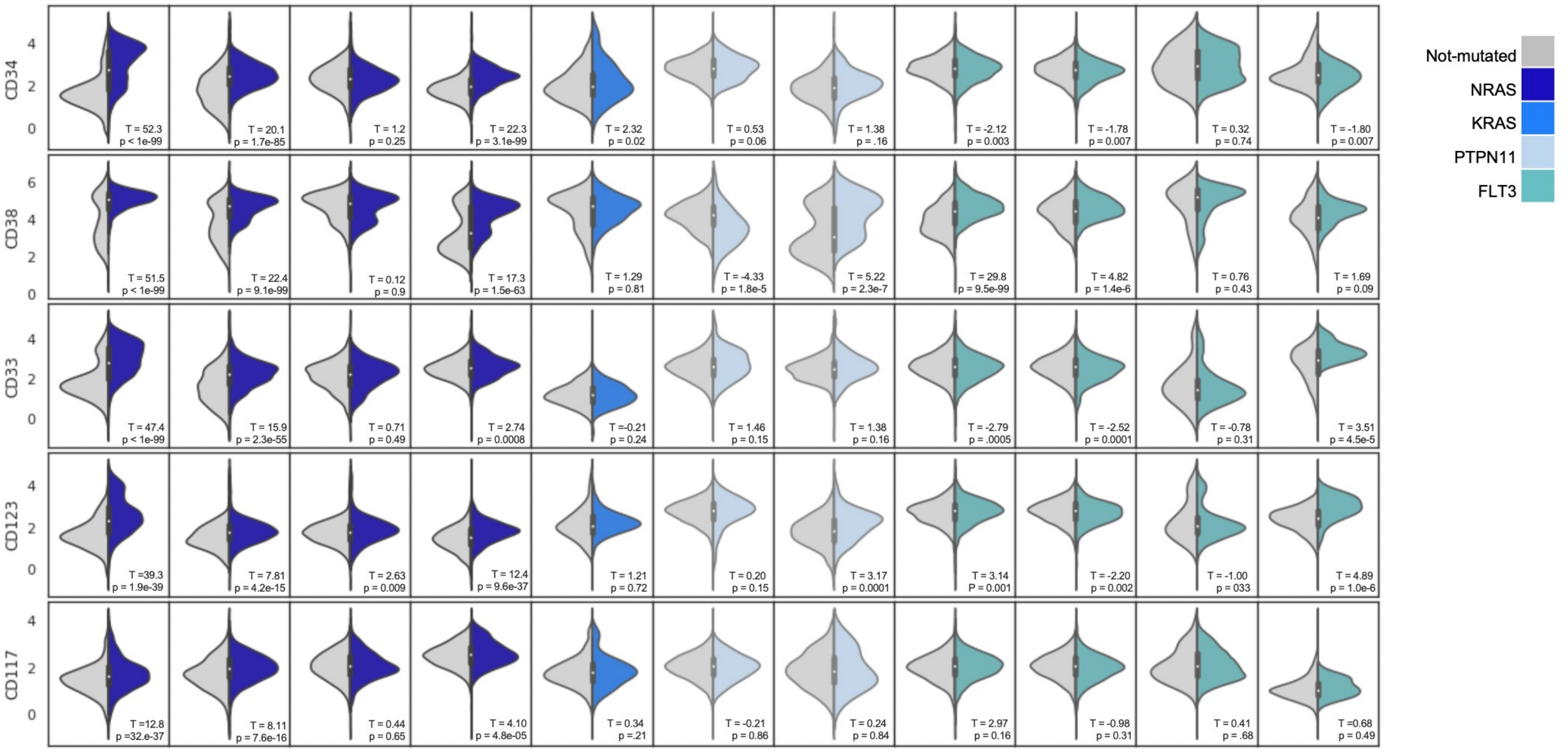


b

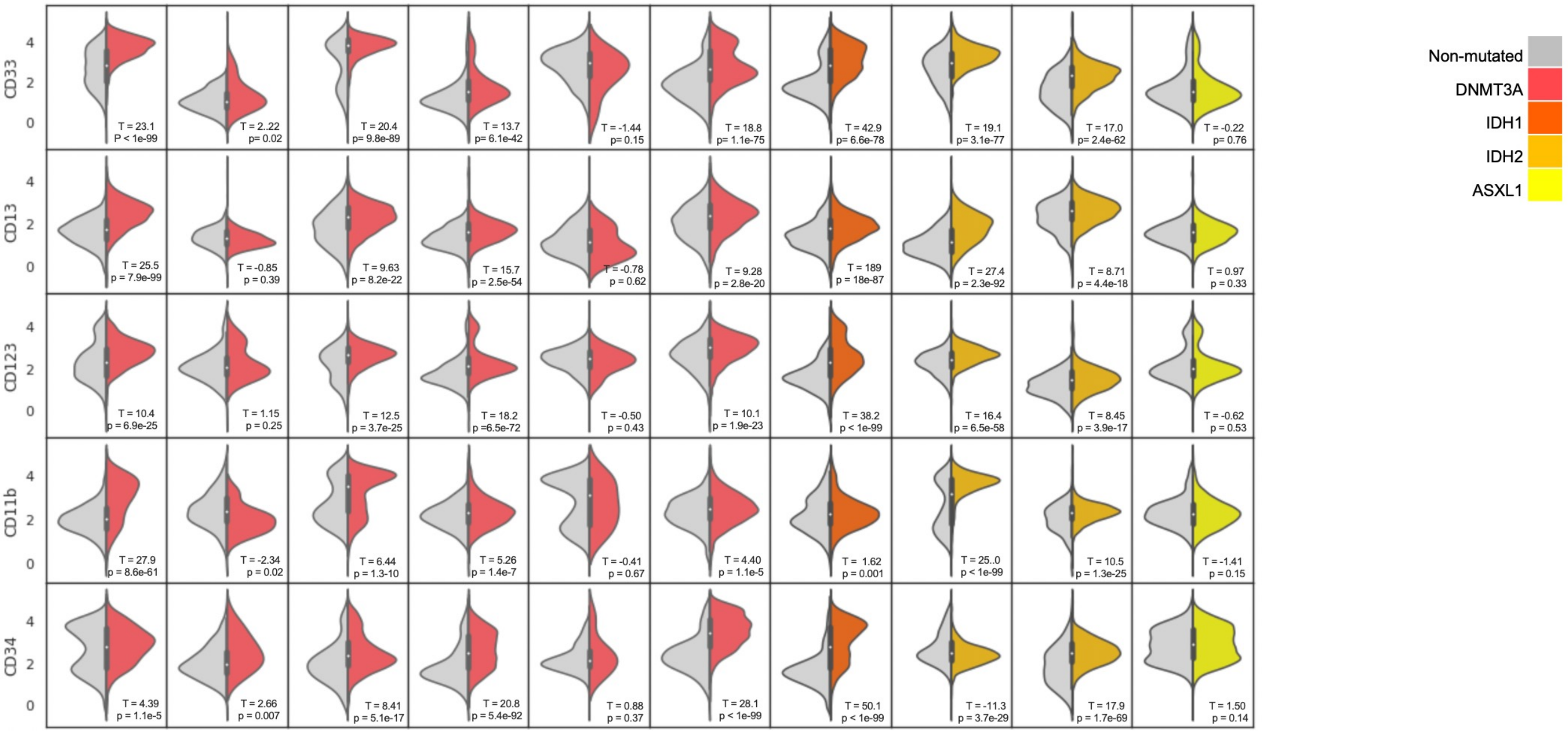


**Supplementary Figure 5.** Heatmap comparing distributions of antibody expression for mutant vs wildtype cells. **A.** Comparison of distributions of antibody expression for mutant vs wildtype cells across 11 population with signaling mutations (NRAS, KRAS, PTPN11, or FLT3). Each column represents a unique mutated population. Each row represents expression of 5 cell surface antibodies with the greatest median T-statistic across all 11 populations (CD34, CD38, CD33, CD123, CD117). The grey half of the split-violin plot represents non-mutated cells and the colorful half of the plot represent mutated cells within an individual patient. **B.** Comparison of distributions of antibody expression for mutant vs wildtype cells across 10 population with epigenetic modifier mutations (DNMT3A, IDH1, IDH2, ASXL1). Each column represents a unique mutated population. Each row represents the expression of 5 cell surface antibodies with the greatest median T-statistic across all 11 populations (CD33, CD13, CD123, CD11b,CD34). The gray half of the split-violin plot represents non-mutated cells and the colorful half of the plot represent mutated cells within an individual patient. All p-values were adjusted via the Bonferroni method for multiple comparisons.

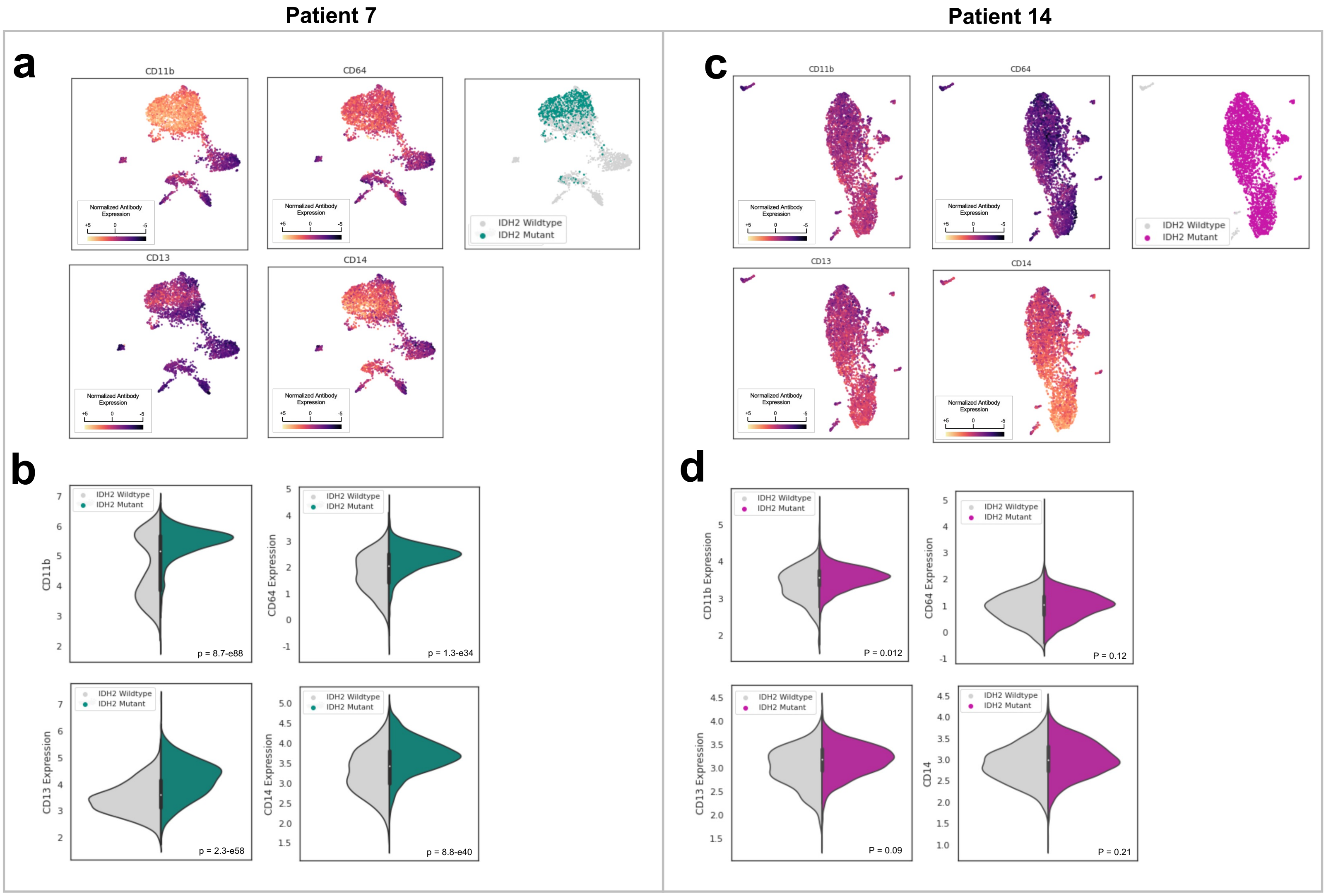


**Supplementary Figure 6.** A. Immunophenotype-derived UMAP from 3,339 cells from Patient 7. Cells are color-coded based on CD11b expression (top left), CD13 expression (bottom left), CD64 expression (top center), CD14 expression (bottom center), and the presence of an IDH2 R140Q mutation (right). B. Violin plot comparing expression of CD11b (top left),CD13 (bottom left), CD64 (top right), and CD14 (bottom right) between IDH2-mutated vs IDH2-wiltype cells in Patient 7. C. Immunophenotype-derived UMAP from 4,082 cells from Patient 14. Cells are color-coded based on CD11b expression (top left), CD13 expression (bottom left), CD64 expression (top center), CD14 expression (bottom center), and the presence of an IDH2 R140Q mutation (right). D Violin plot comparing expression of CD11b (top left),CD13 (bottom left), CD64 (top right), and CD14 (bottom right) between IDH2-mutated vs IDH2-wiltype cells in Patient 4. Statistical significance reflects t-tests and is denoted as *p , .05; **p , .01; ***p , .001, will all p-values adjusted via the Bonferroni method.

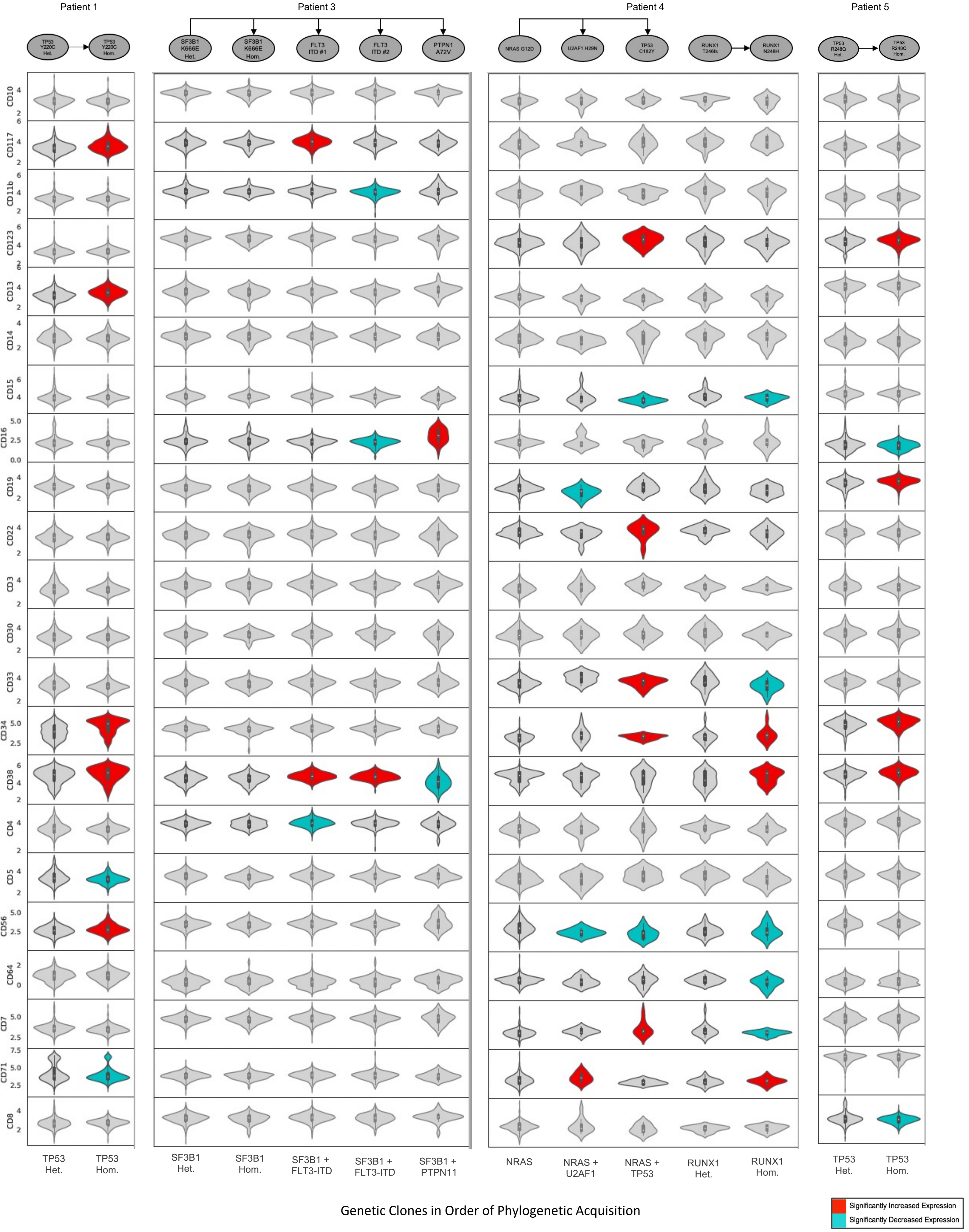

Patient 1
Patient 3
Patient 4
Patient 5
TP53 Y220C Het.
TP53 Y220C Hom.
SF3B1 K666E Het.
SF3B1 K666E Hom.
FLT3 ITD #1
FLT3 ITD #2
PTPN1 A72V
NRAS G12D
U2AF1 H29N
TP53 C182Y
RUNX1 T246fs
RUNX1 N248H
TP53 R248Q Het.
TP53 R248Q Hom.
CD10
CD117
CD11b
CD123
CD13
CD14
CD15
CD16
CD19
CD22
CD3
CD30
CD33
CD34
CD38
CD4
CD5
CD56
CD64
CD7
CD71
CD8
TP53 Het.
TP53 Hom.
SF3B1 Het.
SF3B1 Hom.
SF3B1 + FLT3-ITD
SF3B1 + FLT3-ITD
SF3B1 + PTPN11
NRAS
NRAS + U2AF1
NRAS + TP53
RUNX1 Het.
RUNX1 Hom.
TP53 Het.
TP53 Hom.
Genetic Clones in Order of Phylogenetic Acquisition
Significantly Increased Expression
Significantly Decreased Expression

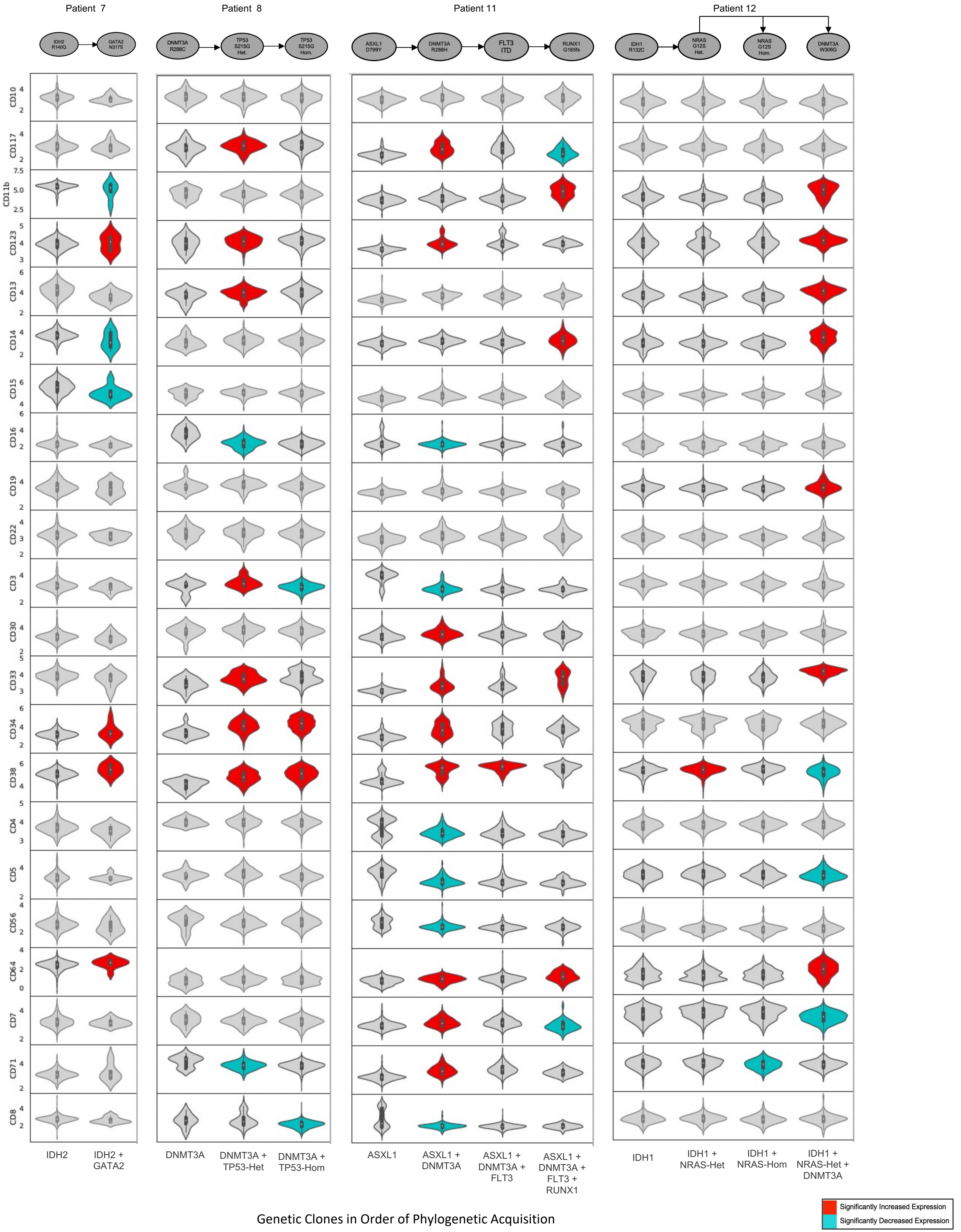

Patient 7
Patient 8
Patient 11
Patient 12
IDH2 R140Q
GATA2 N317S
DNMT3A R288C
TP53 S215G Het.
TP53 S215G Hom.
ASXL1 D799Y
DNMT3A R288H
FLT3 ITD
RUNX1 G165fs
IDH1 R132C
NRAS G12S Het.
NRAS G12S Hom.
DNMT3A W306G
CD10
CD117
CD11b
CD123
CD13
CD14
CD15
CD16
CD19
CD22
CD3
CD30
CD33
CD34
CD38
CD4
CD5
CD56
CD64
CD7
CD71
CD8
IDH2
IDH2 + GATA2
DNMT3A
DNMT3A + TP53-Het
DNMT3A + TP53-Hom
ASXL1
ASXL1 + DNMT3A
ASXL1 + DNMT3A + FLT3
ASXL1 + DNMT3A + FLT3 + RUNX1
IDH1
IDH1 + NRAS-Het
IDH1 + NRAS-Hom
IDH1 + NRAS-Het + DNMT3A
Genetic Clones in Order of Phylogenetic Acquisition
Significantly Increased Expression
Significantly Decreased Expression

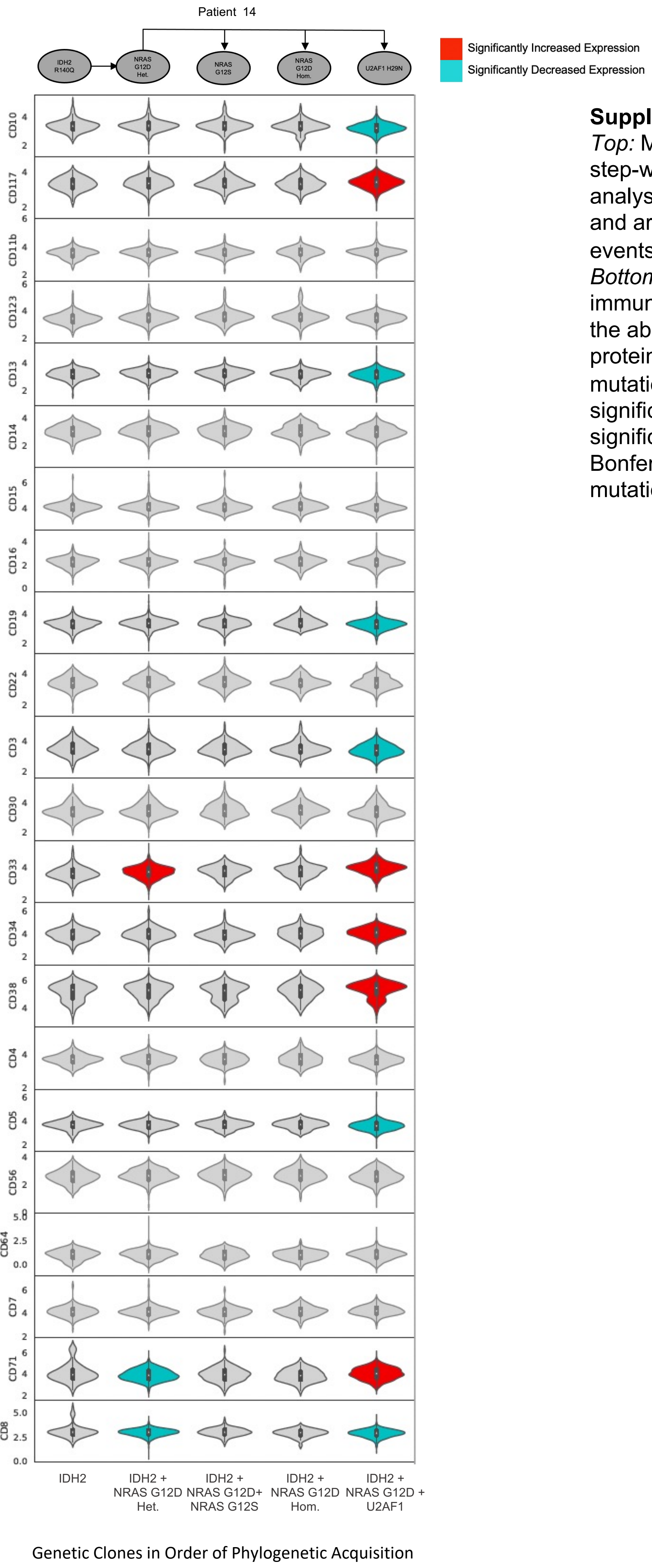


**Supplementary Figure 7.**
*Top:* Mutation phylogeny of 9 patients with MPAL with at least 2 step-wise mutational acquisitions identified on single-cell DNA analysis. Each oval represents a genetically-distinct subclone and arrows represent cumulative acquisition of mutational events.
*Bottom:* Violin plots depicting expression of 22 immunophenotypic proteins for each subclone represented in the above phylogeny. Violin plots color-coded in red indicate protein expression that has significantly increased with mutational acquisition; plots color-coded in blue indicate a significant decrease in protein-expression. Statistical significance is considered $p < 0.05$ after adjustment via the Bonferroni method. Het: Heterozygous; Hom: Homozygous. All mutations are heterozygous unless specified otherwise.

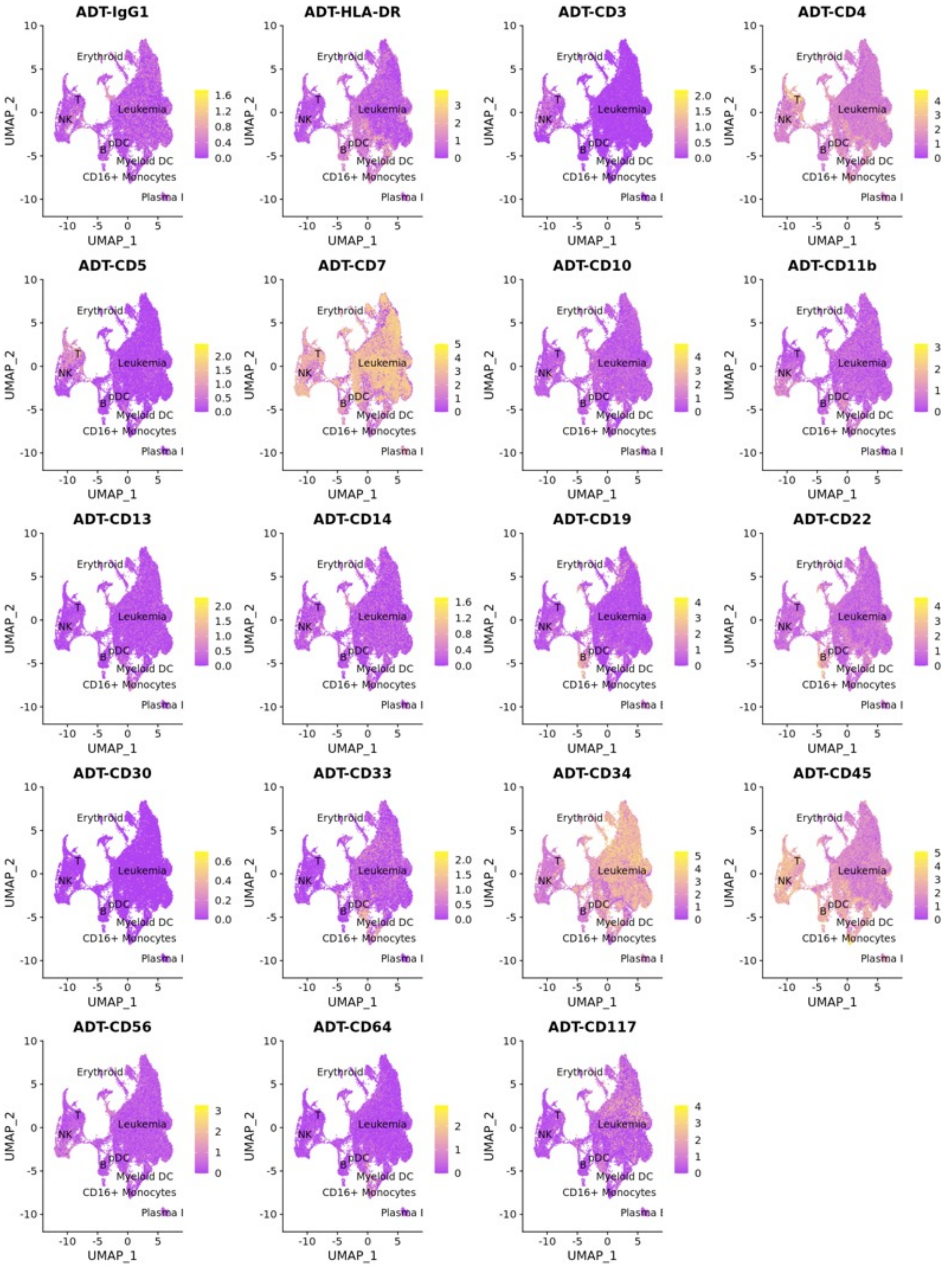


**Supplementary Figure 8.** RNA-derived UMAP from single-cell RNA+protein analysis of 71,579 cells from 12 patients with MPAL. Cells are annotated based on transcriptional data using a combination of scType and clustifyr, with immature populations collapsed into a common 'leukemia' cluster.. Cells are color-coded based on expression of 19 cell-surface proteins.

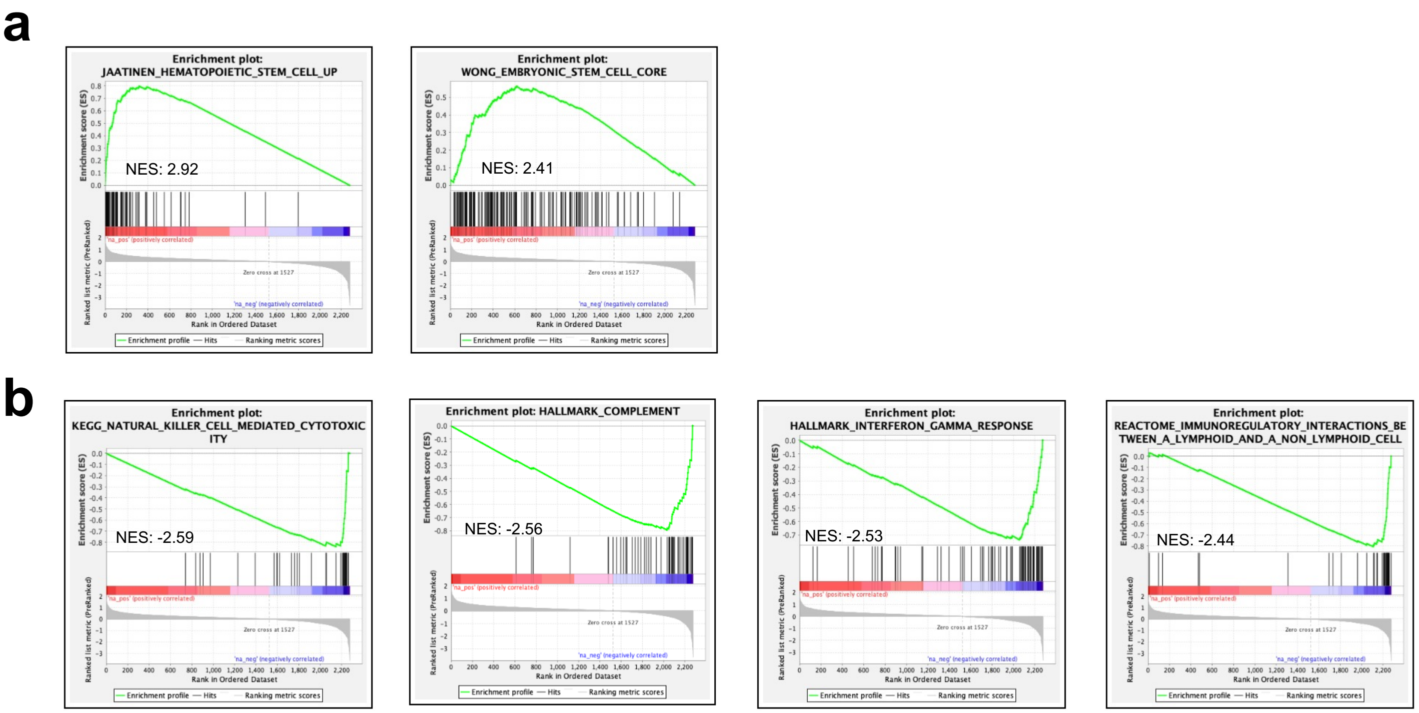


**Supplementary Figure 9. Gene Set Expression Analysis (GSEA) of all cells transcriptionally annotated as Leukemia vs non-Leukemia across 12 patients with MPAL**. A. Enrichment profile and ranking metric score for two example positively enriched gene sets, both associated with stem cells. B. Enrichment profile and ranking metric score for four example negatively enriched gene sets, all associated with immune signaling.

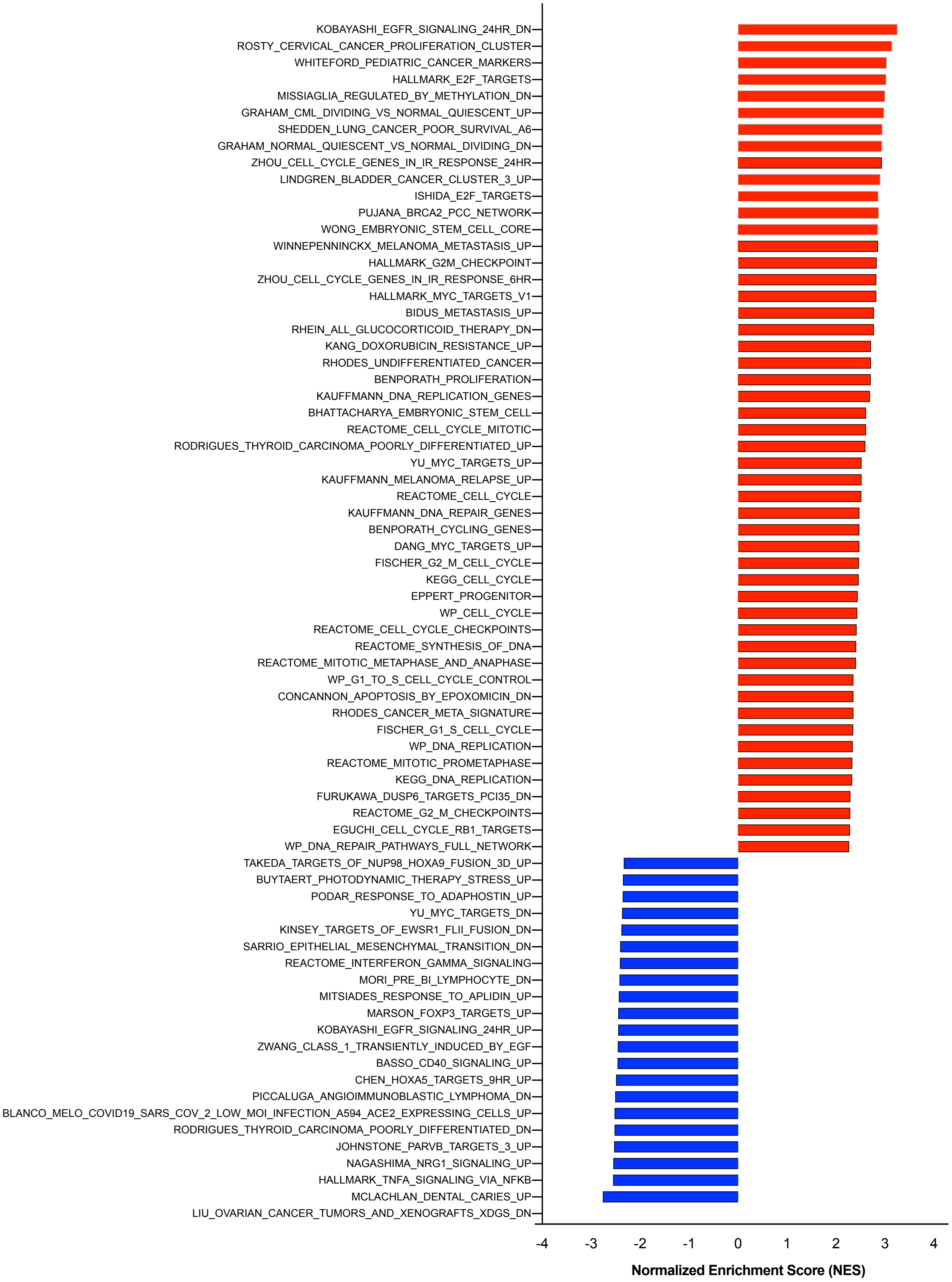


**Supplementary Figure 10. Gene Set Expression Analysis (GSEA) of all leukemia cells with CytoTRACE > 0.95 vs < 0.95**. Bar plot of normalized enrichment score (NES) of 72 gene sets with false-discovery rate q-values < 0.00005. Positively enriched gene sets are color-coded in red and negatively enriched gene sets are color coded in blue.

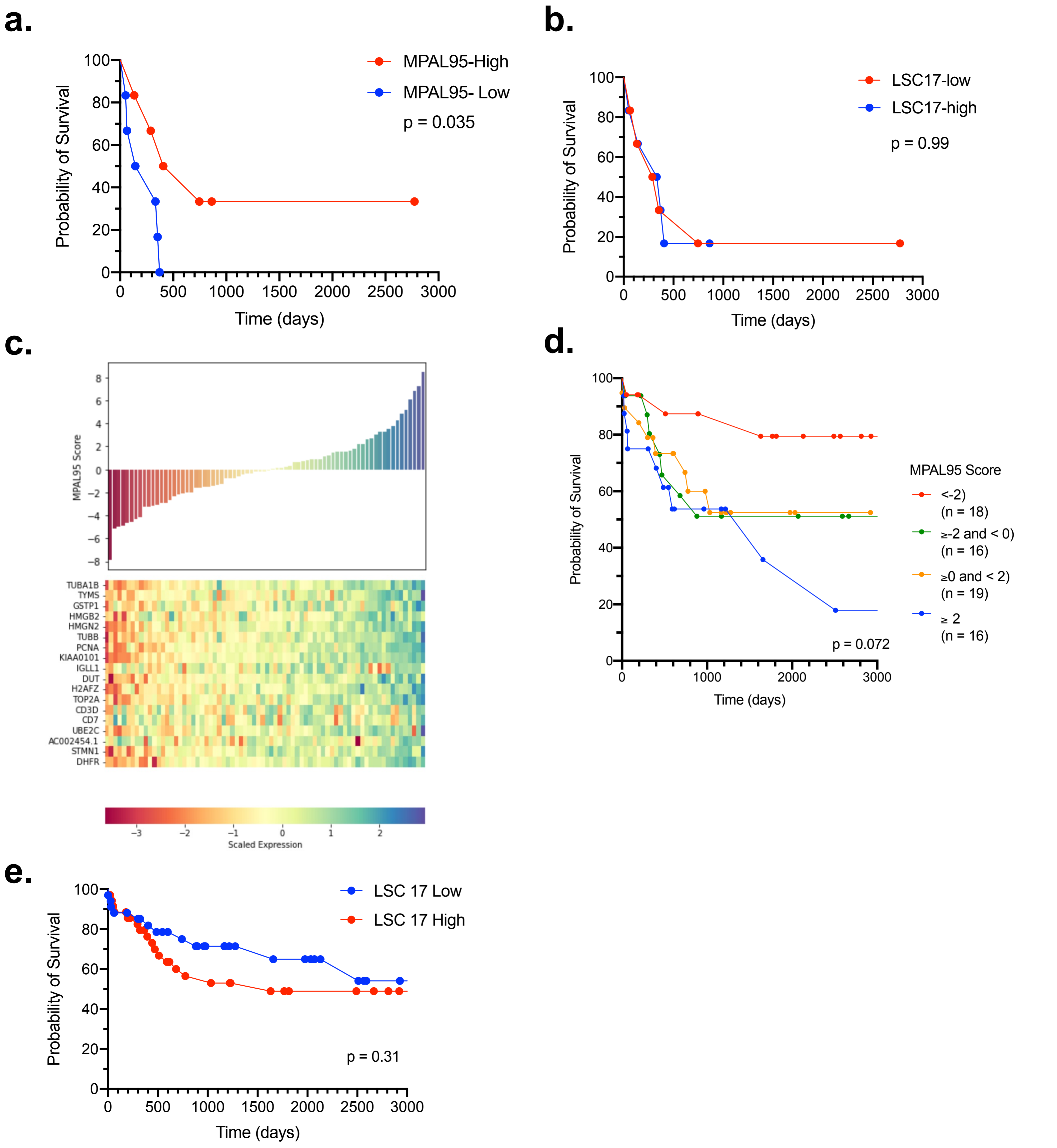


**Supplementary Figure 11.**

A. Kaplan Meier estimates of overall survival stratified by MPAL95 scores for pseudobulked RNAseq data from 12 adult patients with MPAL B. Kaplan Meier estimates of overall survival stratified by LSC17 scores for pseudobulked RNAseq data from 12 adult patients with MPAL C. We generated a gene set score, MPAL95, based on single-data CytoTRACE data and applied it to 69 pediatric patients with survival outcomes available from the TARGET-ALL-P3 dataset. MPAL95 gene set score was computed as the first principal component (top bar plot) of the 18 genes with greatest upregulation in single cells with CytoTRACE scores ≥0.95 (bottom heatmap), where columns are 69 pediatric patients. D. Kaplan Meier estimates of overall survival stratified by MPAL95 scores for TARGET-ALL-P3 data. E. Kaplan Meier estimates of overall survival stratified by leukemia stem cell (LSC) 17 scores for the TARGET-ALL-P3 data.

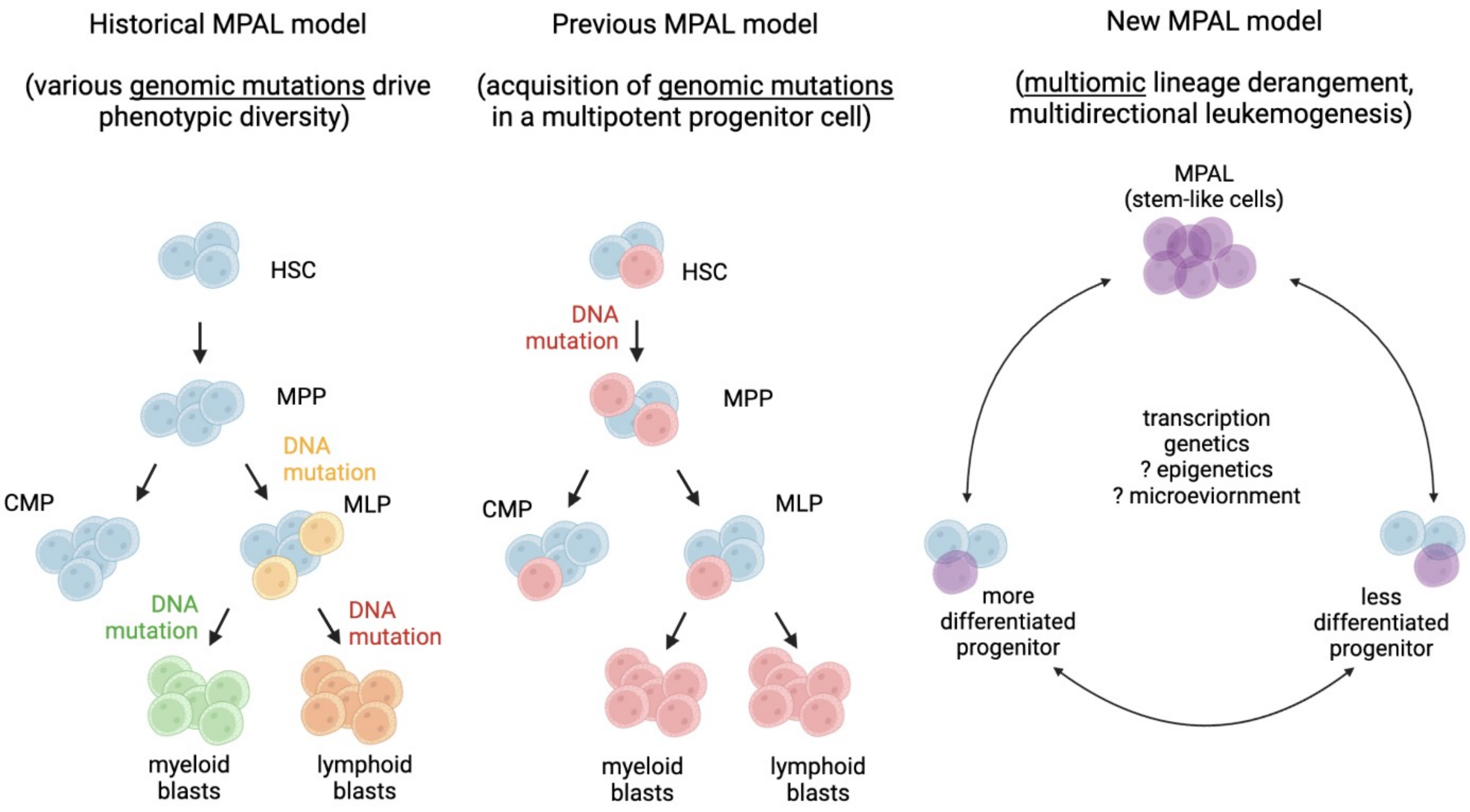


**Supplementary Figure 12. Conclusions from multiomic SC analysis of MPAL**

Potential models of MPAL leukemogenesis. *Left:* Historical model assumed serial acquisition of secondary mutations (yellow, green, orange cells) drove phenotypic diversion, which has been disproven. *Center:* Current model of MPAL assumed acquisition of a driver mutation in a multi-potent hematopoietic progenitor (red cells) that retained lymphoid and myeloid potential so that the inciting mutation could be propagated in different phenotypes. *Right:* Our data suggests that cells may become more phenotypically stem-like with mutational acquisition, and that MPAL is the result of multiomic influence on differentiation potential and phenotypic lineage derangement.

HSC = hematopoietic stem cell; MPP = multipotent progenitors; CMP = common myeloid progenitor; MLP = multi-lymphoid progenitor.
